# Genomic insights into *Ceratobasidium* sp. associated with vascular streak dieback of woody ornamentals in the United States using a metagenomic sequencing approach

**DOI:** 10.1101/2025.08.11.669776

**Authors:** Kassaye H. Belay, Sahar Abdelrazek, Sehgeet Kaur, Reza Mazloom, Devin Bily, Tashi Gyatso, Farhat A. Avin, John Bonkowski, Prabha Liyanapathiranage, Lina Rodriguez Salamanca, Lenwood S. Heath, Fulya Baysal-Gurel, Boris A. Vinatzer

## Abstract

Woody ornamentals are integral to urban landscapes and play important roles in habitat restoration and ecological conservation, yet their national and international trade facilitates the spread of plant diseases with significant ecological and economic consequences. Vascular streak dieback (VSD) recently emerged on woody ornamentals in the United States and was found to be associated with the fungal pathogen *Ceratobasidium sp.* (*Csp*), but little is known about its genomic diversity and associated microbial communities. We thus applied metagenomic sequencing to 106 symptomatic samples that had tested positive for *Csp* and had been collected from 34 woody ornamental species in seven states. Taxonomic profiling identified *Csp* as the only putative pathogen of which we recovered 17 high-quality draft genomes. Phylogenomic and pangenome analyses revealed that U.S. *Csp* isolates form a tight genetic cluster, distinct in gene content from *C. theobromae*, a pathogen of cacao, avocado, and cassava in Southeast Asia. Comparative analyses highlighted gene content differences, including candidate effectors and secondary metabolite clusters, which may underlie host interactions and offer diagnostic targets. These findings provide the first genomic insights into the U.S. *Csp* population, suggest the recent introduction of a single genetic lineage with a broad host range, and establish a framework for improved detection, monitoring, and management of VSD in woody ornamentals.

**Importance:** Identification of the pathogen that causes an emerging disease, be it of humans, animals or plants, is a prerequisite to develop effective treatment and/or management practices and to try to control the disease outbreak to prevent further pathogen spread. Vascular Streak Dieback (VSD) is an emerging disease of ornamental bushes and trees in the U.S. Identification of the pathogen has been hindered by the difficulty in growing the fungal pathogen found to be associated with diseased plants in pure culture. Here we succeeded in sequencing DNA of the likely pathogen directly from plant tissue or from the fungal mass growing out of collected plant tissue. The sequences were assembled into genomes, which allowed us to precisely identify the pathogen, compare it to related pathogens of other plants, and to predict how it causes disease. These results can now be used to inform management and control of VSD.

## Introduction

Plant pathogenic fungi in the genus *Ceratobasidium* are known to cause severe vascular diseases in Southeast Asia, notably vascular streak dieback (VSD) in cacao (*Theobroma cacao*) and avocado (*Persea americana*) (Keane et al., 1972; Guest & Keane, 2007) and witches’ broom disease in cassava (*Manihot esculenta*) (Leiva et al., 2023). Vascular streak dieback (VSD) is characterized by progressive dieback of shoots and branches, often accompanied by leaf yellowing, marginal scorching, and premature defoliation. Infected trees typically show vascular discoloration extending longitudinally in the xylem, which manifests as reddish-brown streaks when stems are split open (Guest et al., 2007; Samuels et al., 2012; McMahon et al., 2016). As the disease advances, plants exhibit stunting, wilt, and canopy thinning, and severely affected trees may experience whole-branch dieback and eventual mortality (Samuels et al., 2012; McMahon et al., 2016). Recently, VSD has also been reported in woody ornamental species across the United States. A *Ceratobasidium* sp. (*Csp*) was first detected in eastern redbud (*Cercis canadensis* L.) with VSD symptoms in Tennessee in 2019, while similar symptoms were observed in maple (*Acer* spp.) in North Carolina as early as 2018 (McClellan, 2023; Liyanapathiranage et al., 2024). To date, the disease has been reported in over 12 states and 50 woody plant genera, with the highest incidence in eastern redbud (*Cercis canadensis* L.), followed by maple (*Acer* spp.) and dogwood (*Cornus* spp.) (Lauderdale, 2023; Baysal-Gurel & Liyanapathiranage; 2024, Bily et al., 2025; Liyanapathiranage et al., 2024). Woody ornamentals play a crucial role in landscaping, habitat restoration, and ecological conservation, and their cultivation and sale significantly contribute to the economic viability of the U.S. nursery industry. The continued spread of VSD is thus a major concern from both an ecological and an economic perspective (Liyanage et al., 2025).

*Ceratobasidium* sp. detection in the U.S. is currently based on a PCR assay that uses primers specific to the *Csp* ITS region (Avin et al., 2025). Phylogenetic reconstruction has been performed using the ITS, LSU, ATP6, TEF1, and RPB2 genes and revealed a close genetic relationship between *Csp* in the U.S. and *C. theobromae* (*Ct*) from cacao and cassava in Southeast Asia but since *Csp* sequences form a distinct clade, they may represent a separate, previously unrecognized species (Liyanapathiranage et al., 2024). Also, the 100% sequence identity in the ITS region between the U.S. *Csp* sequences and *Ct* sequences from Japanese honeysuckle (*Lonicera japonica*) in China suggest that *Csp* in the U.S may be more closely related to *Ct* in China than to *Ct* in Southeast Asia (Liyanapathiranage et al., 2024). Koch’s postulates have not been completed because of the fastidious nature of *Csp*. Mycelium grows out of plant tissue but, so far, is recalcitrant to sub-culturing. Therefore, it cannot be excluded that other biotic or abiotic factors contribute to the disease.

The genera *Ceratobasidium* and *Rhizoctonia* are closely related basidiomycete fungi within the family *Ceratobasidiaceae*, exhibiting overlapping morphological and ecological traits that complicate their differentiation (Moliszewska et al., 2023). Historically, many *Ceratobasidium* species were identified as the sexual (teleomorphic) stage of *Rhizoctonia* (González García et al., 2006; Gónzalez et al., 2016). Recent phylogenetic investigations have further blurred genus boundaries, revealing that some species formerly classified under *Ceratobasidium* are more accurately placed within *Rhizoctonia* (Moliszewska et al., 2023; Gónzalez et al., 2016). Regarding pathogenicity, *Rhizoctonia* is notorious for causing a broad range of diseases such as damping-off and root rot (Moliszewska et al., 2023). In contrast, *Ceratobasidium* species generally adopt a more specialized biotrophic lifestyle, maintaining a more intimate and often less aggressive interaction with their hosts (Keane et al., 1972; Guest & Keane, 2007).

Genomic studies of *Ct* have greatly advanced understanding of this pathogen’s biology and evolution (Ali et al., 2019; Gil-Ordóñez et al., 2024). These studies revealed compact genome sizes of approximately 33–34 Mb encoding 9,000–10,000 genes. Comparative genomic analyses identified around 800 genes unique to *Ct* compared to its closest relatives potentially underpinning *Ct*’s vascular pathogenicity, including an extensive secretome, plant cell wall degrading enzymes, and virulence-associated factors such as toxin biosynthesis and effector proteins (Ali et al., 2019; Gil-Ordóñez et al., 2024).

A comprehensive understanding of the emergence, distribution, transmission, and evolutionary dynamics of emerging plant diseases, such as VSD, is essential for developing effective disease containment or management strategies (Rasmussen and Grünwald, 2021; Woolhouse et al., 2005). Today, for culturable pathogens, this information can be obtained by whole genome sequencing (WGS) with long-read platforms, like PacBio and Oxford Nanopore Technologies, which facilitate accurate assembly and characterization of complex fungal genomes (Wang et al., 2021; Sun et al., 2022). However, WGS becomes challenging or impossible to use in the case of fastidious or non-culturable pathogens, like *Csp*. Metagenomic sequencing, *i.e.*, the direct sequencing and analysis of all genetic material obtained from an environmental or host sample without requiring cultivation (Handelsman et al., 2004), can potentially provide the same information as WGS. It can even identify mixed infections. But metagenomics is faced with several challenges. In the case of plant samples, overabundance of host sequences limits the number of pathogen sequences that can be obtained. Also, the presence of many other microbes may make it challenging to separate pathogen sequences from non-pathogen sequences, which limits the possibility to obtain a complete pathogen genome with minimal contamination. Therefore, metagenome-assembled genomes (MAGs) of plant pathogens have so far mainly been obtained for bacteria (Mande et al., 2012; Meyer et al., 2022; Johnson et al., 2022; Roman-Reyna and Crandall., 2024; Abdelrazek et al., 2025).

To start answering the open questions about VSD on woody ornamentals in the U.S., here, we sequenced 106 samples from plants with VSD symptoms that had been confirmed to be infected with *Csp* with a PCR assay (Liyanapathiranage et al., 2024; Avin et al., 2025), including 61 symptomatic vascular tissue samples and 45 mycelia from 34 different woody plant species collected in seven U.S. states. The main objectives were to (i) perform a survey of bacterial, fungal, and oomycete pathogens present in symptomatic vascular tissues to identify the microbial diversity in VSD symptomatic tissue, including the detection of *Csp* and other co-existing microorganisms, (ii) assemble *Csp* genomes from plant metagenomic and mycelium samples to elucidate evolutionary relationships among *Csp* sequences from the U.S. and with *Ct* sequences from Southeast Asia, and (iii) assess differences in gene content, including predicted virulence genes, between *Csp* and *Ct* to start unraveling potential adaptation of *Csp* to woody ornamentals and the various regional climates in North America.

## Methods

### Plant Sample Collection

Survey sites included wholesale and retail nurseries, along with landscape sites with newly planted trees. Symptomatic native plants located near landscape trees or nursery stock at each survey site were also sampled. Sampling was performed as described previously (Bily et al., 2025). In short, one to three plant samples were chosen from a block, group, or cultivar of plants that best represented the symptoms observed, which included marginal leaf scorching, leaf chlorosis, stunting, wilt, vascular discoloration, and dieback. These symptoms were consistently observed across all samples at each site. Symptomatic plant parts or the entire plant with roots were placed in sealed plastic bags at room temperature and were either shipped overnight or delivered the same day to the laboratory. Fungal cankering and oomycete root rot pathogens were occasionally associated with symptomatic plants. However, note that only plants that tested positive for *Csp* were used further (see next section). **Table S1** provides comprehensive information including host plant species, geographic collection locations, and isolation dates.

### Plant Sample Processing, culturing, DNA Extraction, and PCR

Samples were processed as previously described (Bily et al., 2025). In short, after removing foliage, stems were cut into 15 to 20 cm pieces, surface sterilized in 70% ethanol, and incubated at ≤90% humidity at 22 ± 2°C in the dark for 24 to 96 h. Using a scalpel, 0.1 g of discolored xylem tissue was cut out lengthwise or sliced into discs from smaller, debarked shoots. If parts of the plant were dead, any discolored living tissue at the margin between dead or wilted parts was selected. If the sample lacked dead tissue or vascular discoloration, four pieces (0.025g each) of xylem tissue would be selected every 5 cm and combined, making sure to select from multiple growth rings. Tissue was then macerated in CD1 buffer of the Qiagen DNeasy Plant Pro Kit (Germantown, MD) with a BioSpec MiniBeadbeater-16 (Bartlesville, OK) or MP BiomedicaFastPrep-24™ 5G (Solon, OH), followed by DNA extraction using the manufacturer’s protocol. Conventional PCR and qPCR targeting *Csp* was used on extracted DNA as previously described (Avin et al., 2025; Bily et al., 2025).

If stems had sufficient moisture content, culturing from vascular tissue was attempted, where a 4 to 5 mm piece of discolored xylem tissue was surface sterilized with 2.5% NaOCl, again with 70% ethanol, and air-dried before being placed on water agar medium (pH 7.1) and incubated in the dark at 22 ± 2°C for as long as 15 d, where 0.1 g of *Rhizoctonia*-like mycelia was swabbed with a sterile needle into an extraction tube with CD1 buffer. For other stem samples, mycelium was harvested under a dissection microscope with a sterile needle directly from cut stems incubated for 96 h. DNA was then extracted from mycelium using again the Qiagen DNeasy Plant Pro Kit.

### Computational Resources

All bioinformatic analyses were performed on the OWL’s Nest and TinkerCliffs high-performance computing clusters (Advanced Research Computing, ARC) at Virginia Tech. Analyses typically used 20–40 CPUs and up to 256 GB of RAM per job, and all major software tools and pipelines were installed and managed through centrally maintained conda environments.

### Metagenomic sequencing

Long-read sequencing was performed on an Oxford Nanopore Technologies (ONT) PromethION sequencer. Sequencing libraries were prepared following the manufacturer’s protocol for SQK-NBD114.24, provided by ONT. Briefly, nucleic acid extracted from each sample went through an FFPE repair step and then purified using AMPure XP beads (Beckman Coulter Life Sciences, Indianapolis, IN). Following purification, sequencing adapters were ligated to each sample, then subjected to a final purification step using AMPure XP beads. For each sequencing library that was prepared, barcoded and adapter-ligated samples were pooled together and eluted in 25 µl of ONT elution buffer. PromethION flow cells (FLO-PRO114M) were primed, and the sequencing library concentration and quality were assessed using Qubit and TapeStation, then adjusted to meet PromethION requirements before being loaded into the primed flow cells for sequencing. The resulting FASTQ files from each flow cell were concatenated into a single file per sample for downstream analysis. All reads that passed ONT quality control (fastq_pass) were utilized in subsequent analyses. ONT reads were basecalled using Guppy v6.3.7 in high-accuracy mode, and reads were quality filtered with NanoFilt v2.8.0 using a minimum average Phred quality score of Q ≥ 10 and a minimum read length of 1,000 bp. Reads not meeting these thresholds were discarded. After concatenation, each FASTQ file was evaluated to determine read statistics, including the total number of reads, total base count (the sum of all read lengths), and the average read length (**Table S1**). These metrics were calculated using a custom Bash script that parsed the FASTQ files by counting sequence lines and summing their lengths to generate the summary statistics.

### Taxonomic Profiling

To identify plant host sequences, the read mapping tool Minimap2 (Li, 2018) was used, choosing as reference genomes the plant genome assembly available in NCBI that was most closely related to the host of isolation (accession numbers are listed in **Table S1**). Minimap2 was also used to identify *Csp* sequences by mapping reads against the two available *Ct* genomes from cacao: CT2 (accession GCA_009078325.1) and Gudang 4 (accession GCA_012932095.1). Reads that mapped against both genomes were de-duplicated before downstream analysis.

The taxonomic profiler Kraken2 (Breitwieser et al., 2018; Wood et al., 2019; Lu et al., 2023) was employed to assign sequencing reads to bacterial, fungal, and oomycete taxa. Two databases were used for identification of bacterial and fungal taxa, respectively: (1) the standard k2_pluspfp_20240605 database, comprising RefSeq archaea, bacteria, viral, plasmid, human, protozoa, fungi, plants, and UniVec_Core (Langmead, 2024), and (2) a custom fungi-oomycete database to address the limited representation of these taxa in RefSeq. To generate the custom database, 18,644 fungal and 513 oomycete genome assemblies were downloaded from NCBI on July 24, 2024. These assemblies were processed using LINflow (Tian et al., 2021) to assign Life Identification Numbers (LINs). Subsequently, representative genomes were selected at an average nucleotide identity (ANI) threshold of 99.5%, resulting in a non-redundant, comprehensive Kraken2 database containing 8,797 fungal and 342 oomycete genomes.

Additionally, taxonomic profiling were performed using sourmash v4.5.0 (Brown and Irber, 2016; Irber et al., 2024). Raw sequencing reads were converted into FracMinHash sketches with a k-mer size of 31 and scaled parameter of 1000. Bacterial community composition was determined using the sourmash `gather` command against the GTDB-RS207 reference database. To analyze fungal communities, the genomes used above for the fungi-oomycete Kraken2 database were used to build a custom sourmash database with the same FracMinHash parameters as for bacteria.

### *Csp* Genome Assembly and Analysis

Samples containing more than 10,000 reads mapping to the two *Ct* reference genomes (GCA_009078325.1 and GCA_012932095.1) were selected for genome assembly. Each selected sample was assembled independently using Flye (Kolmogorov et al., 2019), specifying a genome size parameter of 34 Mb based on the *Ct* reference genome size. Five rounds of iterative polishing were conducted to enhance assembly accuracy.

To capture *Csp*-specific genomic regions potentially missed by aligning exclusively to *Ct* genomes, we constructed a comprehensive assembly by pooling the five metagenomic samples containing the highest abundance of *Ct*-specific reads. Host-derived sequences were first removed using Minimap2. The remaining non-host reads were combined and subjected to *de novo* assembly using Flye (Kolmogorov et al., 2019). Following this assembly, initial contig filtering was conducted using Kraken2 with both standard and custom fungi-oomycete databases. Contigs were further analyzed using BLASTn searches (Altschul et al., 1990) against the NCBI core nucleotide database with relaxed alignment parameters, maximizing detection of divergent sequences. Additionally, read-depth consistency analyses were performed on each contig using Samtools (Li et al., 2009), CoverM (Aroney et al., 2025), and mosdepth (Pedersen et al., 2017). Only contigs displaying high sequence similarity (≥95% nucleotide identity over ≥1 kb) to non-*Ceratobasidiaceae* taxa, or those exhibiting inconsistent read-depth patterns, were excluded. This approach minimized the risk of inadvertently removing genuine Csp sequences.

### Phylogenetic Analysis

For core-genome construction, genome completeness was first assessed using BUSCO v5.2.2 with the Agaricomycetes_odb10 dataset, comprising 2,898 single-copy orthologs (Manni et al., 2021). Conserved single-copy orthologs present in all genomes used for tree construction were aligned using MUSCLE (Edgar, 2004), and alignments were trimmed with TrimAl (Capella-Gutiérrez et al., 2009) to remove poorly aligned regions that could introduce artifacts in phylogenetic reconstruction. A maximum likelihood tree was then inferred using IQ-TREE (Nguyen et al., 2015). Branch support was evaluated using ultrafast bootstrap approximations implemented in IQ-TREE (v2.2.0.3) (Nguyen et al., 2015) under the VT+F+I+I+R6 model with 1,000 ultrafast bootstrap replicates. The final phylogenetic trees were visualized and annotated in R version 4.2.3 using the ggtree package (Yu et al., 2017).

For the single nucleotide polymorphism (SNP) analysis, raw reads were first host-filtered to remove plant sequences. The remaining non-host reads were aligned to two Ct reference genomes, CT2 (GenBank GCA_014217725.1) and Gudang 4 (GenBank GCA_012932095.1) to extract *Ceratobasidium* (*Csp*) reads. The extracted *Csp* reads were then mapped to the *Ct* reference isolate LAO1 (GenBank GCA_037974915) using minimap2 v2.29 with the `-ax map-ont` preset. Alignments were sorted and indexed with samtools (v1.19.1). Variants were called per isolate using NanoCaller (v3.6.2) in haploid mode (`--mode all, --sequencing ul_ont,--snp_model ONT-HG002, --indel_model ONT-HG002`), producing one VCF containing SNPs and indels per sample. VCFs were coordinated□sorted, BGZF□compressed and indexed with `bcftools` and `tabix`, then merged into a single multi□sample VCF (`bcftools merge --force-samples`). The merged VCF was filtered to retain only high□confidence SNPs (per□site QUAL ≥ 20 and per□sample depth DP ≥ 10) using bcftools `view -i ‘QUAL>=20 && FMT/DP>=`10’. The resulting filtered VCF was converted to a multi□FASTA alignment in PHYLIP format using a custom vcf2phylip script. A maximum□likelihood phylogeny was inferred in IQ-TREE (v2.2.0.3) (Nguyen et al., 2015) under the `GTR+`Γ model with 1,000 ultrafast bootstrap replicates. The final phylogenetic tree was visualized and annotated in R version 4.2.3 using the ggtree package (Yu et al., 2017).

### Gene Prediction and Comparative Genomics

Gene prediction was performed using AUGUSTUS v3.5.0 (Stanke et al., 2006) with an *ab initio* approach trained on the *Rhizoctonia solani AG-1 IA* genome (GenBank: GCF_016906535.1). We trained the classifier on *R. solani* due to its well-curated genome and comprehensive effector predictions, whereas *C. theobromae* currently lacks equivalent functional annotation, limiting its utility for reliable training and benchmarking. The training pipeline began by initializing species-specific configuration files using `new_species.pl` to establish baseline parameters, followed by automated gene model training with `autoAugTrain.pl` utilizing multicore optimization. Pre-processing of annotations retained only mRNA and CDS features to ensure AUGUSTUS training compatibility, resulting in optimized gene model parameters for accurate prediction in *Csp* genomes. Following gene prediction, protein-coding sequences were extracted from GFF3 annotations using `gffread` (Pertea et al., 2020) and underwent quality control curation. OrthoFinder (Emms et al., 2019) performed orthologous clustering using the DIAMOND (Buchfink et al., 2021) algorithm in a multi-threaded configuration, while custom Python scripts generated presence/absence matrices to define core and accessory orthogroups with gene IDs mapped to their respective genomes. Core orthogroup protein sequences were concatenated. The concatenated core protein sequences underwent MAFFT (Kazutaka et al., 2019) alignment using 40 threads for optimal performance, followed by TrimAl processing to remove poorly aligned regions that could interfere with phylogenetic signals. Seqmagick converted the trimmed alignment into PHYLIP format for compatibility with downstream analysis tools. IQ-TREE performed maximum likelihood phylogenetic reconstruction using the `VT+F+I+I+R6` substitution model, with branch support assessed through both ultrafast bootstrap and SH-aLRT (1000 replicates each), ultimately providing evolutionary relationship insights among *Csp* isolates.

### Functional Genome Annotation

Protein sequences from core and accessory genomes were functionally annotated using InterProScan (Quevillon et al., 2005). Before annotation, sequences were quality-checked to retain only valid amino acid residues. InterProScan analyses utilized multiple protein databases (Pfam, SMART, CDD, etc.) to annotate protein domains, GO terms, and metabolic pathways. InterProScan was executed on high-performance computing resources, allocating 150 GB RAM and 16 CPU cores. Outputs included TSV, XML, and GFF3 formats, providing extensive functional annotations suitable for downstream enrichment and comparative analyses. Quick summary tables were generated to count the frequency of InterPro domains, GO terms, and associated pathways, facilitating functional enrichment analyses to identify potential host-specific adaptations and evolutionary innovations in *Csp* genomes.

### Prediction and Characterization of Putative Effector Proteins

Putative effector proteins were identified from the predicted accessory proteome following a multi-step computational pipeline integrating secretion, localization, and functional prediction tools (Logachev et al., 2024; Poudel et al., 2023). Candidate protein sequences derived from accessory orthogroups were first screened for the presence of N-terminal signal peptides using SignalP v6.0 (Teufel et al., 2022) in eukaryotic mode with fast processing settings. Signal peptide-positive sequences were then analyzed with TMHMM v2.0c (Krogh et al., 2001; Käll et al 2007) in short output mode to identify proteins containing transmembrane helices. Proteins predicted with one or more helices were excluded to only retain putative soluble secreted proteins. Proteins lacking transmembrane helices were further screened using TargetP v2.0 (Armenteros et al., 2019) in non-plant eukaryotic mode to identify and exclude proteins predicted to contain mitochondrial targeting peptides (mTP). The remaining secretome candidates were scanned for endoplasmic reticulum retention signals (PS00014 motif) using ScanProsite (de Castro et al., 2006). Proteins carrying this motif were excluded to avoid ER-retained proteins. The refined secretome was finally evaluated for glycosylphosphatidylinositol (GPI) anchor signals using NetGPI v1.1 (Gíslason et al., 2021). Candidate proteins were split into batches and submitted to the web server for prediction. Proteins identified as GPI-anchored were excluded from the secretome. The filtered secretome was submitted to EffectorP v3.0 (Sperschneider & Dodds, 2022) in fungal mode to predict candidate effector proteins. EffectorP provided classification outputs distinguishing effector and non-effector proteins, along with localization categories as apoplastic, cytoplasmic, or dual-localized effectors. EffectorP prediction outputs were parsed to separate candidate effectors by localization class. Protein sequences corresponding to each effector class were extracted using seqkit (Shen et al., 2016) and compiled into dedicated FASTA files.

## Results

### Collection of 106 symptomatic *Csp*-positive samples from 34 plant species in seven U.S. states

Plant tissue specimens exhibiting VSD symptoms were collected from various woody ornamental hosts across seven U.S. states. Sampling criteria included the presence of vascular discoloration, foliar chlorosis progressing to necrosis, and terminal dieback consistent with previously documented VSD symptoms (**Figure 1**). The frequency of symptoms varied with nursery, ranging from a few scattered plants to 90% of the stock. A total of 106 samples wer obtained and tested positive for *Csp* based on a diagnostic PCR assay (Avin et al., 2025). Virginia contributed the largest sample cohort (n=64, 58.2%), followed by Tennessee (n=25, 22.7%), North Carolina (n=7, 6.4%), Indiana (n=4, 3.6%), Texas (n=3, 2.7%), Maryland (n=2, 1.8%), and Missouri (n=1, 0.9%) (**Figure 2A**). Within Virginia, specimens were collected from 14 counties with the greatest concentration in the central Piedmont region (Orange, Louisa, Fauquier, Nelson, Loudoun and Albemarle) (**Figure 2B**), which contains the highest density of commercial woody ornamental nurseries in the state.

**Figure 1.**
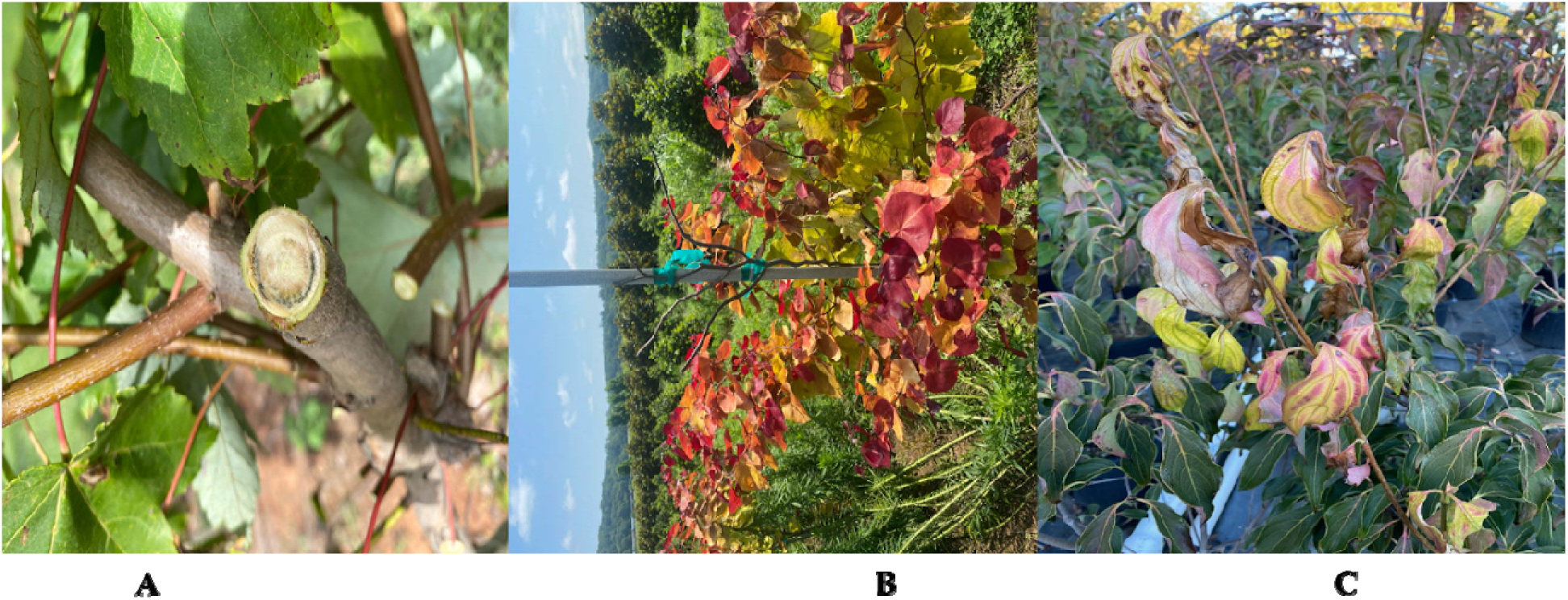
Representative vascular streak dieback (VSD) symptoms observed in *Csp*-positive woody ornamental hosts: (A) *Acer rubrum* ‘October Glory’; (B) *Cercis canadensis* ‘Flame Thrower’; (C) *Cornus kousa* ‘Greensleeves’. Symptoms include vascular discoloration, foliar chlorosis progressing to necrosis, and terminal dieback.

**Figure 2.**
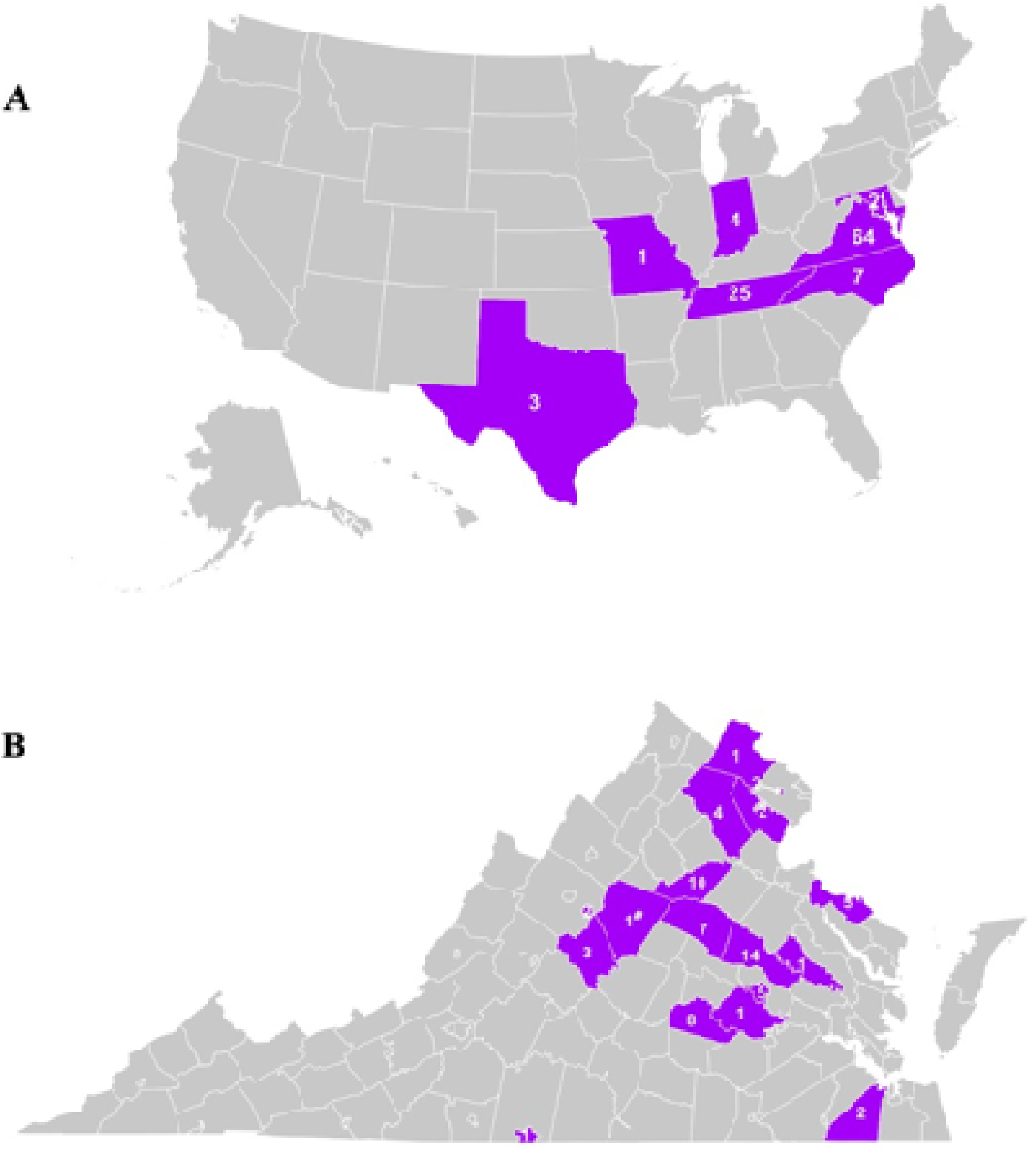
Geographic distribution of *Csp* sample collection sites across the United States. **A** map of the U.S. with the states from which samples were obtained highlighted in purple. The numbers indicate the number of samples collected from each state. **B** A map of Virginia showing counties and number of samples that were collected.

The samples originated from 34 woody ornamental host species belonging to 19 different plant families (**Table S1**), with eastern redbud (*Cercis canadensis*, n=28), Freeman maple (*Acer × freemanii*, n=18), and flowering dogwood (*Cornus florida*, n=16) comprising the majority of specimens. Additional hosts included, but were not limited to, Japanese maple (*Acer palmatum*), southern wax myrtle (*Morella cerifera*), Chinese dogwood (*Cornus kousa*), crepe myrtle (*Lagerstroemia indica*), and various oak species (*Quercus* spp.).

### Deep metagenomic sequencing provides insight into microbial communities associated with VSD

Sequencing was performed on DNA either extracted from vascular tissue (61 samples) or from mycelium cultured directly from vascular tissue (45 samples). Sequencing yielded a cumulative total of 188,856,320 reads across all samples. The sequencing depth per sample was highly variable, ranging from 14,364 to 7,555,390 reads, with an average of 1,760,656 reads (see detailed sequencing results for each sample in **Table S1**). The mean read length for all sequencing runs was 2,940 base pairs (bp) with the longest obtained read being 396,120 bp long, resulting in a combined total of approximately 893 gigabases (Gb) of sequence data. The high sequencing depth in many samples promised to enable comprehensive profiling of microbial communities and robust detection of *Csp* even in samples with low *Csp* abundance.

Not surprisingly, plant reads represented the majority of reads in sequenced plant samples (67.8% on average, with a range from 50.01–99.01%) but plant reads also accounted for a substantial fraction of mycelium derived samples averaging 31.7%, with a range 0.3–88.2% (**Figure 3**). This was likely because mycelium frequently grew intermixed with remnants of vascular plant tissue and small amounts of host material were occasionally co-harvested during sample preparation. As expected, fungal and bacterial read counts were higher in mycelium compared to plant samples (approximately 10-fold) because of the lower average number of plant reads. However, in both sample types, bacterial read and fungal read numbers were similar to each other. Reads assigned to oomycetes were overall much lower, but relatively higher in plant samples compared to mycelium samples.

**Figure 3.**
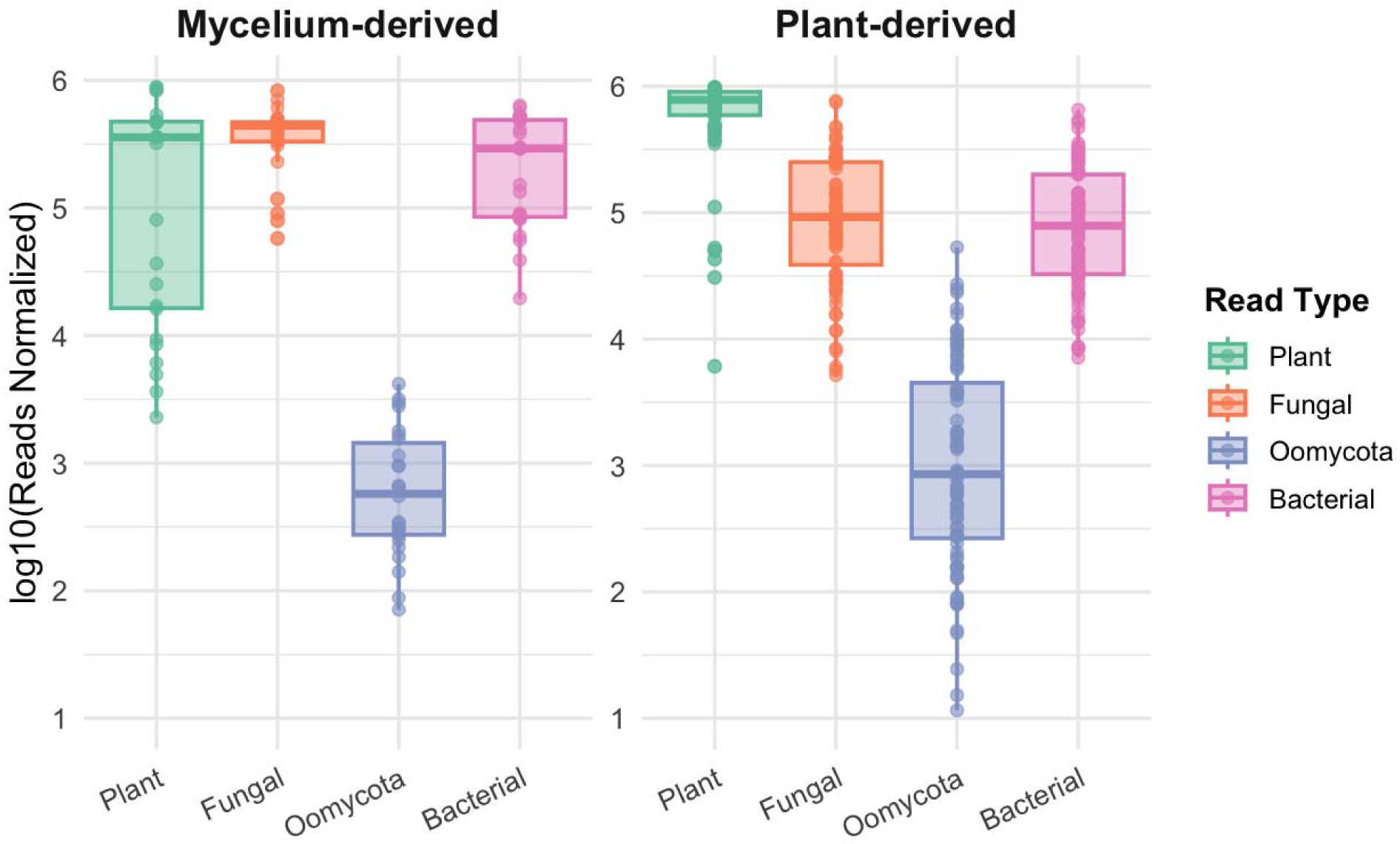
Boxplots of log□□-transformed normalized read counts (reads per million) for plant, fungal, Oomycota, and bacterial taxa, comparing Mycelium-derived (left) and plant-derived (right) samples.

Previous investigations of VSD in woody ornamentals in the U.S. focused on the fungal pathogen *Csp* (Liyanapathiranage et al., 2024, Avin et al., 2025; Bily et al 2025. Here, metagenomic sequencing allowed us to investigate the entire microbial communities associated with VSD and determine if any other microbe consistently co-occurs with *Csp*. Fungal taxonomic classification using the Kraken2 tool (Breitwieser et al., 2018; Wood et al., 2019; Lu et al., 2023) with a custom fungal and oomycete database revealed distinct fungal community patterns between mycelium and plant samples. In mycelium samples, *Csp* was the dominant taxon, ranking first in 23 of the 30 samples with relative abundance from 19.6% to 91.9%, exceeding half of all non-host reads in 20 samples. The number of *Csp* reads was independently confirmed by mapping reads against *Ct* genomes, which largely aligned with the taxonomic profiling results (**Figure 4** and **Table S1**). Based on Kraken2, *Fusarium falciforme*, a pathogen that causes wilt disease and root rots in many herbaceous and some wooden species (Thao et al., 2021; Siddig et al., 2023), was the dominant taxon in the remaining mycelium samples (**Figure 5A**, **Table S1**). However, when mapping these reads against a *F. falciforme* reference genome, on average only 0.3% of reads identified by Kraken2 as *F. falciforme* mapped to *F. falciforme* genomes. When these reads were assembled into contigs, the contigs could not be confirmed by BLAST searches against a custom *Fusarium* database. Further validation using the taxonomic profiler sourmash (Brown and Irber, 2016; Irber et al., 2024) with a custom fungi and oomycete database containing the same genomes used for the Kraken2 database also failed to detect *F. falciforme* (**Table S2**).

**Figure 4.**
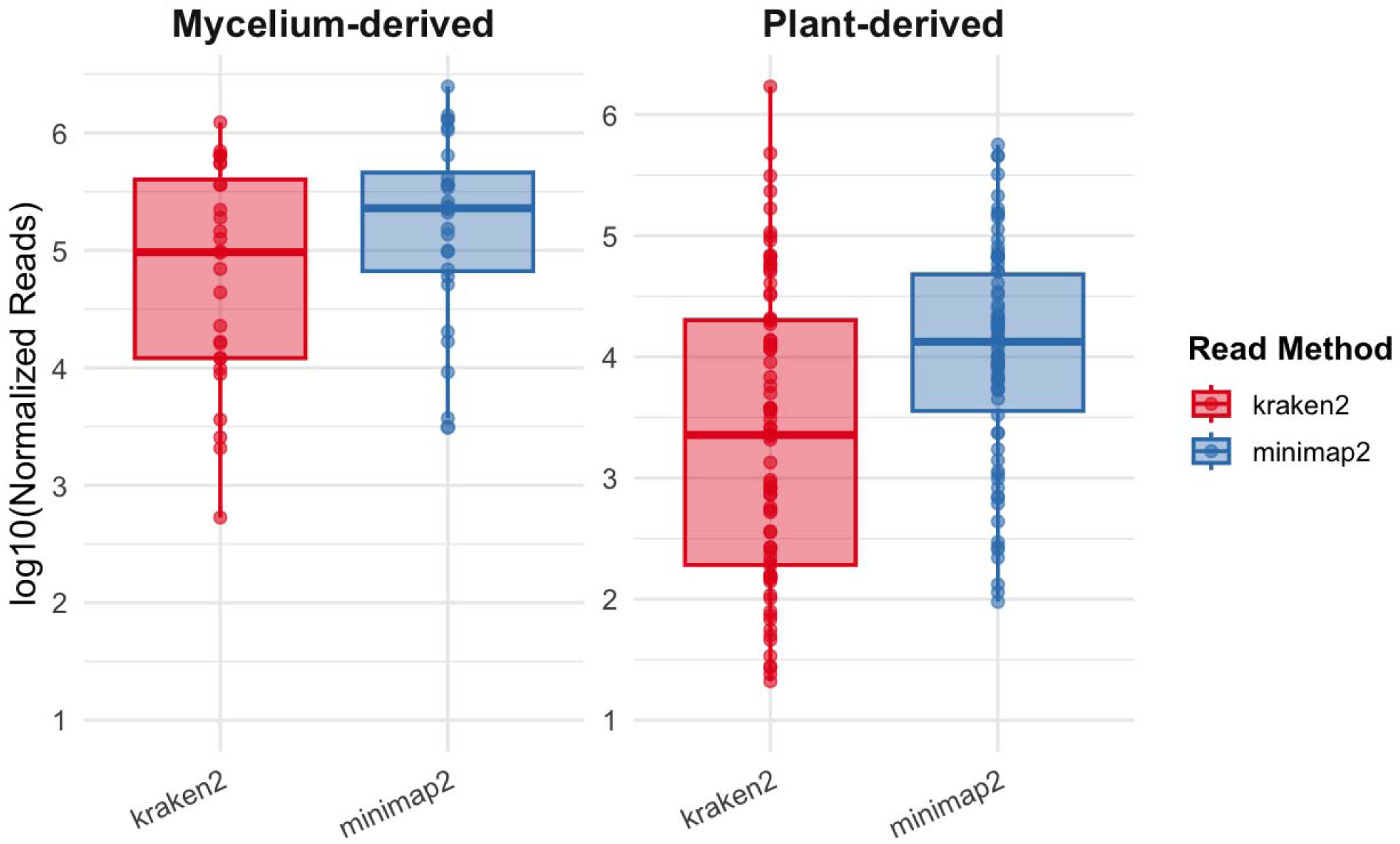
Boxplots of log□□-transformed normalized read counts of *Csp* comparing Kraken2 (red) and Minimap2 (blue) across mycelium-derived (left) and plant-derived (right) samples. In both sample types, Minimap2 shows a slightly higher median and tighter distribution of read counts than Kraken2.

**Figure 5.**
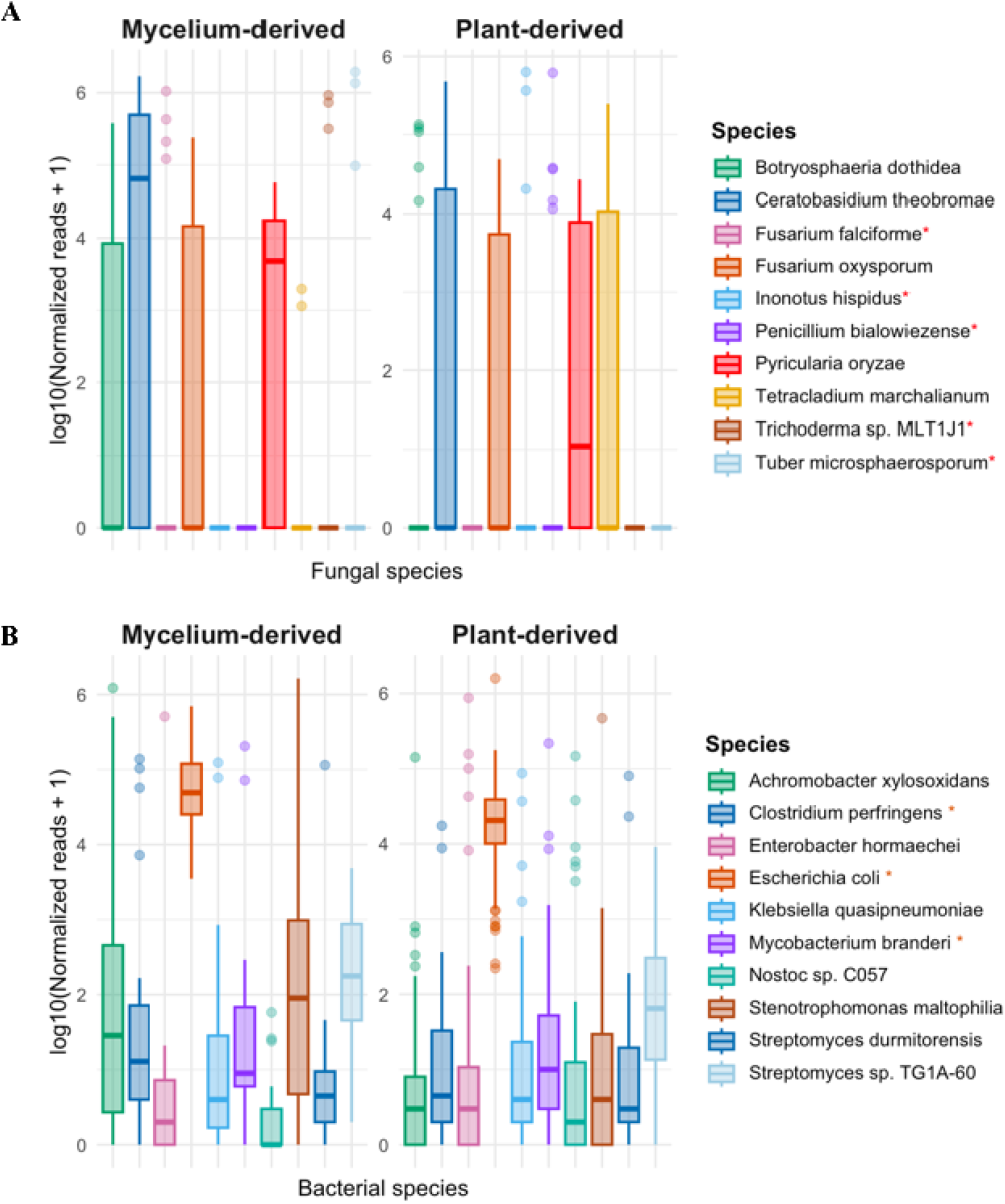
Two-panel boxplots of log□□-transformed normalized reads for the top 10 taxa identified by Kraken2. **(A)** Fungal species and **(B)** Bacterial species, each comparing mycelium-derived (left) and plant-derived (right) samples. Boxes show the interquartile range with the median (thick line); whiskers extend to 1.5×IQR; points are per-sample values. Species marked with an asterisk (*) in the legends were not supported by read-mapping or sourmash and are therefore likely Kraken2 false positives.

Plant samples exhibited greater variability regarding the most highly abundant fungal species than mycelium samples (**Figure 5A**, **Table S1**). Only in 16 of 76 plant samples was *Csp* the most abundant fungus with relative abundances ranging from 6.33% to 82.46% out of all non-host reads. As in mycelium samples, when *Csp* was not dominant, *F. falciforme* was often the most dominant species based on Kraken2 (but, as for mycelium samples, these reads could not be confirmed). *Botryosphaeria dothidea*, a canker and dieback pathogen of trees with a broad host range (Marsberg et al., 2017) and previously excluded from causing VSD (Liyanapathiranage et al., 2024), was found to be abundant in 13 plant samples. Although *B. dothidea* was confirmed by both sourmash and read mapping against a *B. dothidea* reference genome, when attempting assembly of *B. dothidea* reads, only highly fragmented short assemblies were obtained suggesting lower abundance than that inferred by Kraken2 (**Table S2**). *Pyricularia oryzae* was another species frequently found among the most abundant taxa, ranking within the top three in 58 plant samples. The presence of *P. oryzae* was confirmed using the taxonomic profiler sourmash but reads extracted by mapping to a reference genome could not be assembled into any contigs indicating an overestimation of abundance, and likely misclassification, by Kraken2 and sourmash (**Table S2**). Other fungal species detected by Kraken2 included the endophyte *Tetracladium marchalianum*, the broad host-range pathogen *Fusarium oxysporum*, the environmental fungus *Penicillium bialowiezense*, the powdery mildew pathogen *Erysiphe pulchra*, the mushroom *Psilocybe semilanceata*, and the endophyte (and occasionally pathogen) *Trichoderma spp* (**Figure 5A**, **Table S1**). However, none of the pathogens in this list were present with enough consistency and sufficient abundance to suggest them as contributing to VSD. Moreover, follow up analyses did not confirm their presence at the levels of abundance that Kraken2 determined (**Table S2**).

Oomycetes represented a minor component of the non-host reads in both sample types based on Kraken2 using our custom fungi-oomycetes database accounting on average for only 1.08% of all non-host reads in plant samples and for only 0.36% in mycelium samples. *Plasmopara halstedii* and *Plasmopara obducens* were the most frequently detected species across both sample types (**Table S1**). Other regularly detected oomycetes included *Pseudoperonospora cubensis* and various *Phytophthora* species. We thus conclude that oomycetes are unlikely to play a role in VSD.

Metagenomic analysis of bacteria revealed thirty-seven genera among the top three most abundant bacterial taxa across samples (**Table S1**). Relative abundances varied between plant samples and mycelium samples. In plant samples, members of the *Enterobacteriaceae* including *Enterobacter hormaechei*, *Escherichia coli*, and *Klebsiella quasipneumoniae* were among the most prevalent, while mycelium-derived samples were characterized by higher relative abundances of *Stenotrophomonas maltophilia, Clostridium perfringens*, and *Achromobacter xylosoxidans* (**Figure 5B**, **Table S1**). To validate these taxonomic assignments, we performed read mapping against reference genomes for the five most abundant bacterial species. Of these, only assignments to *Achromobacter xylosoxidans* and *Stenotrophomonas maltophilia* were confirmed and enough reads were found to assemble the respective genomes, whose identities were confirmed using genomeRxiv (Pritchard et al., 2022). For *Escherichia coli*, results varied significantly across samples, prompting additional verification using the FracMinHash-based sourmash tool, which indicated that most reads assigned to *E. coli* by Kraken2 more closely matched genomes of various environmental and commensal taxa, such as *Methylobacterium* sp. *Bradyrhizobium sp.*, and *Curtobacterium* sp. suggesting potential misclassification by Kraken2. Likewise, reads assigned by Kraken2 to *Clostridium perfringens* and *Mycobacterium branderi* aligned to other taxa in the sourmash analysis (**Table S3**). In summary, no bacterial pathogen was consistently identified across samples strongly suggesting that bacterial pathogens do not contribute to VSD.

### Genome Assemblies of *Ceratobasidium species* (*Csp*) isolates

*Csp* reads identified by mapping against the two *Ct* reference genomes were used to construct metagenome-assembled genomes (MAGs). Mapped reads were extracted, deduplicated, and assembled. High-quality draft genome assemblies were obtained from 17 samples (3 from plant samples and 14 from mycelium samples), comparable in size and completeness to the *Ct* reference genomes based on BUSCO completeness scores (**Table 1**). Comprehensive statistics including read counts, genome completeness, genome sizes, and BUSCO scores are detailed in **Table S4**. Twenty-three other samples yielded partial genome assemblies due to lower read counts (fewer than 59,918 reads) with genomes sizes ranging from 2.3 to 30.2, while 46 samples had such a small number of *Csp* reads (fewer than 14,177) that no meaningful assemblies were obtained.

**Table 1:** Summary statistics of *Csp* genome assemblies and *Ct* reference genomes.

| Sample ID | Assembly Length (Mb) | N50 (bp) | No. of Contigs | Annotated Genes | No. of Singletons | BUSCO (Agaricomycetes_odb10) | BUSCO (Fungi_odb10) | Sample Type |
| --- | --- | --- | --- | --- | --- | --- | --- | --- |
| 23_02068_EasternRedbud_IN | 32.97 | 21,383 | 2,325 | 10,442 | 356 | 72.7 | 83.3 | PD<sup>1</sup> |
| FBG5639_EasternRedbud_TN | 36.94 | 53,266 | 2,040 | 11,655 | 567 | 82.0 | 91.9 | PD |
| RT23_468_FloweringDogwood_VA | 38.11 | 28,656 | 2,614 | 10,971 | 320 | 78.1 | 89.3 | PD |
| RT23_372_FloweringDogwood_VA | 36.84 | 161,947 | 786 | 10,990 | 35 | 83.6 | 94.2 | MD<sup>2</sup> |
| RT23_384_FloweringDogwood_VA | 36.78 | 460,389 | 562 | 10,936 | 8 | 84.0 | 94.6 | MD |
| RT23_392_SouthernCatalpa_VA | 36.86 | 549,311 | 483 | 10,779 | 2 | 83.7 | 94.5 | MD |
| RT23_402_RedOsierDogwood_VA | 36.01 | 643,614 | 475 | 10,887 | 0 | 83.9 | 94.6 | MD |
| RT23_423_ChineseDogwood_VA | 36.65 | 390,318 | 543 | 10,678 | 9 | 83.9 | 94.6 | MD |
| RT23_424_ChineseDogwood_VA | 35.37 | 139,030 | 915 | 10,919 | 38 | 83.6 | 94.2 | MD |
| RT23_428_ChineseDogwood_VA | 37.00 | 616,318 | 539 | 10,939 | 1 | 83.8 | 94.5 | MD |
| RT23_429_ChineseDogwood_VA | 36.86 | 700,228 | 485 | 10,880 | 3 | 83.9 | 94.5 | MD |
| RT23_433_FloweringDogwood_VA | 36.53 | 816,925 | 401 | 11,141 | 5 | 83.9 | 94.6 | MD |
| VSD24_268_Sourwood_VA | 32.89 | 37,465 | 1,409 | 10,217 | 311 | 72.2 | 82.7 | MD |
| VSD24_269_RedOsierDogwood_VA | 36.25 | 266,961 | 559 | 10,863 | 24 | 83.6 | 94.6 | MD |
| VSD24_270_FloweringDogwood_VA | 36.11 | 97,071 | 669 | 10,913 | 177 | 80.4 | 91.7 | MD |
| VSD24_DSB_110M_SouthernWaxMyrtle_VA | 36.82 | 379,450 | 293 | 10,779 | 28 | 83.6 | 94.6 | MD |
| VSVD24_368_EasternRedbud_VA | 36.18 | 320,509 | 504 | 10,792 | 13 | 83.9 | 94.6 | MD |
| Comprehensive_assembly | 37.75 | 824,000 | 518 | 10,962 | 5 | 83.7 | 94.3 | Mixed |
| *Ct* CT2 | 31.20 | 86,100 | 1,240 | 9,763 | 104 | 83.6 | 94.5 | NCBI |
| *Ct* Gudang 4 | 38.60 | 39,200 | 4,661 | 13,311 | 1,201 | 82.0 | 92.2 | NCBI |

<sup>1</sup> mycelium-derived (MD)519 <sup>2</sup> Plant-derived (MD)

Since construction of the *Csp* MAGs relied exclusively on reads mapped to the *Ct* reference genomes, we were concerned that we may have failed to capture genomic regions potentially present in *Csp* but absent from *Ct*. Therefore, we conducted a second, independent assembly approach to include these potential regions unique to *Csp*. To do this, we selected the five samples with the highest *Ct*-aligned read counts. Instead of assembling mapped reads, we removed host sequences and then assembled all other reads (2,520,850 reads with a total length of 12.1Gb). The assembly resulted in 522 contigs. We then taxonomically classified these contigs using BLASTn and Kraken2 and removed 4 contigs that had best hits to non-fungal organisms (**Table S5**). While detailed assembly results are shown in **Figure S1**, **Table 1** shows that the comprehensive assembly is very similar in quality to the best individual assemblies. Importantly, the number of annotated genes in the comprehensive assembly (10,962) fell within the range of annotated genes in the individual assemblies (10,442 to 11,141) and, based on our pangenomes analysis (see more details below), only five singletons were found in the comprehensive assembly suggesting that the best individual *Csp* assemblies were not missing a significant number of genes.

### Phylogeny to resolve evolutionary relationships between *Csp* samples from woody ornamentals in the U.S. and related species and genera

To resolve the evolutionary relationships between *Csp* genomes from the U.S., other publicly available *Csp* and *Ct* genomes, and related *R. solani* genomes, a core phylogenetic tree was constructed based on 76 conserved single-copy orthologs. Only the 13 U.S. *Csp* genome assemblies with the highest BUSCO scores were used. The resulting maximum likelihood tree (**Figure 6A**) revealed a well-supported monophyletic clade composed exclusively of U.S. *Csp* isolates. The relationships among most of these isolates remains unresolved with bootstrap support below 50, leading to a striking polytomy. The Southeast Asian *Ct* genomes (CT2 and Gudang4 from cacao) formed a separate, well-supported clade distinct from the U.S. *Csp* lineage. The distinct branch of the single LAO1 genome from cassava suggests that it may be the only representative of a yet undersampled additional *Ceratobasidium* clade. These genome all shared a most recent common ancestor distinct from a sister clade of all other *Ceratobasidium* species. Three *R. solani* genomes, chosen to root the tree, formed yet another, well-separated clade. In summary, the core genome tree confirms that the *Csp* isolates from woody species in the U.S. only recently diverged from a common ancestor and that they are closely related to, but distinct from, the *Ct* isolates from cacao and cassava in Southeast Asia.

**Figure 6A.**
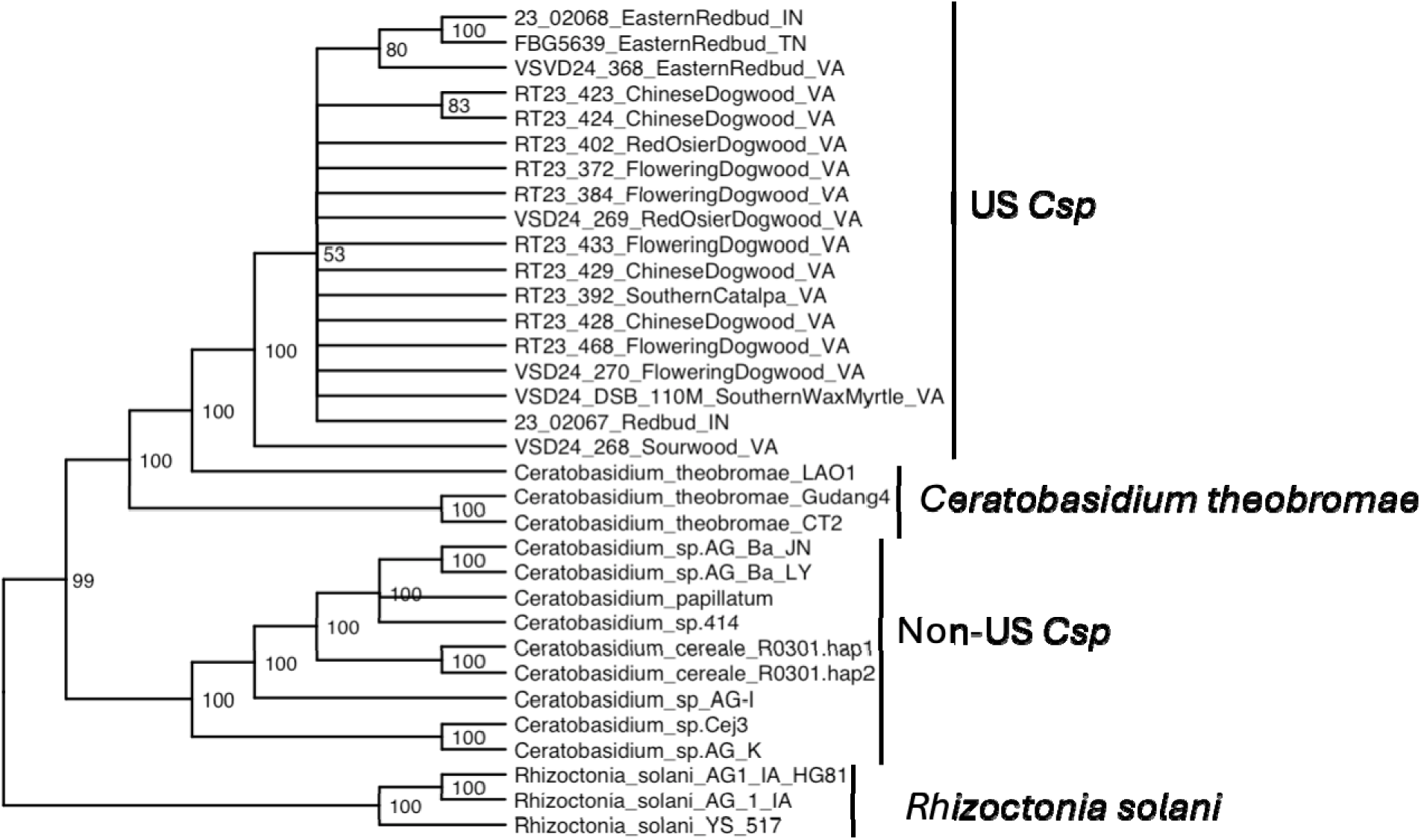
Maximum likelihood phylogenetic tree constructed using 76 single-copy orthologous genes from assembled U.S. *Csp* genomes, publicly available *Ceratobasidium* genomes, and *R. solani* isolates used as an outgroup to root the tree (best model: `VT+F+I+I+R6`). Branch tips are labeled with the sample IDs. Bootstrap support values are shown at each node.

**Figure 6B:**
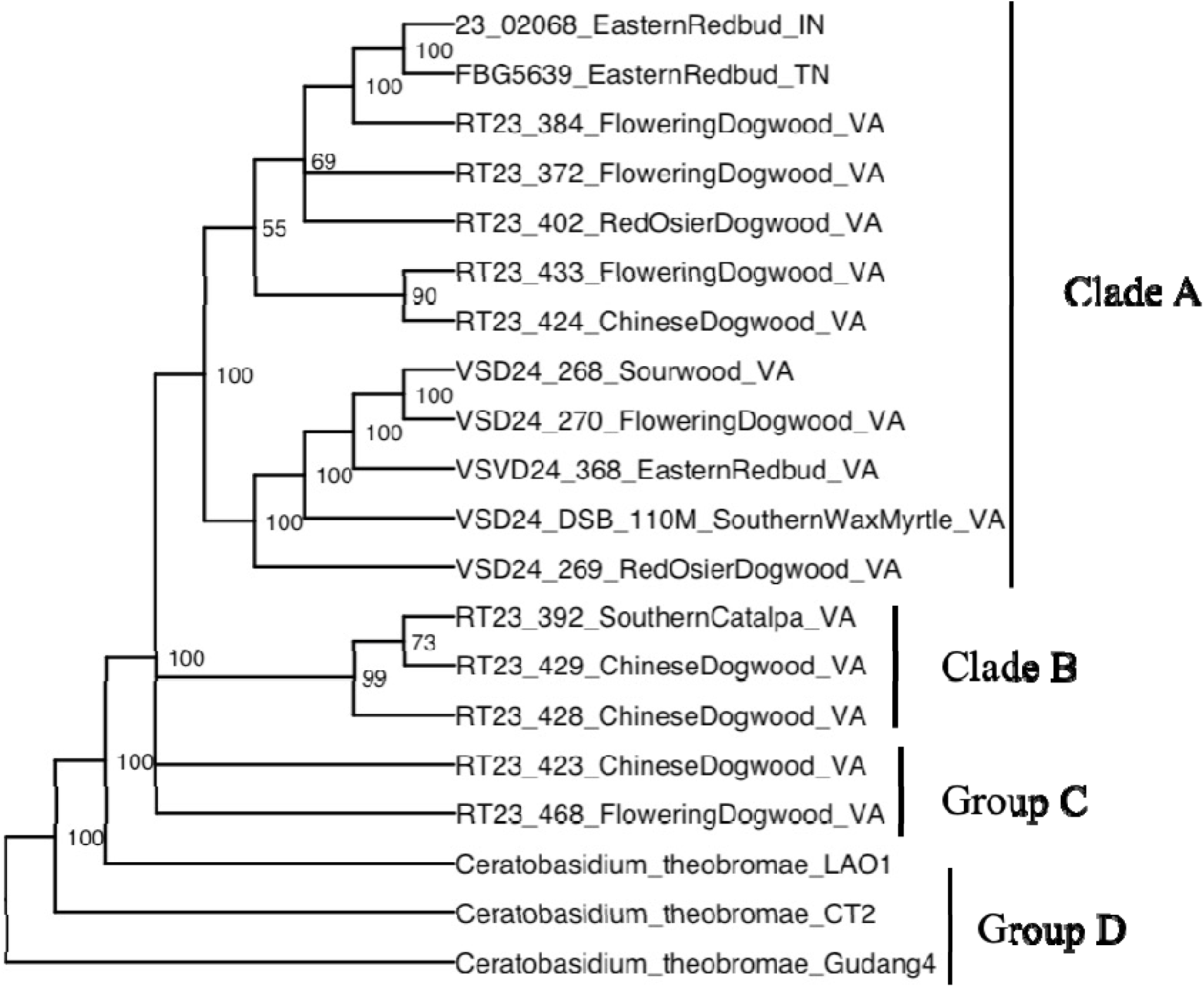
Core genome phylogenetic tree of *Csp* genomes recovered from woody ornamental hosts in the United States (best model: `VT+F+I+I+R6`). A total of 1,473 core genes were used to construct the tree, which was rooted on genomes of *Ct* isolates from cacao and cassava in Southeast Asia. Branch tips are labeled with the sample IDs.

### Phylogeny to resolve evolutionary relationships among *Csp* samples from woody ornamentals in the U.S

A finer-scale analysis focusing on the 17 U.S. *Csp* assemblies of the highest quality and th three publicly available *Ct* genomes based on 1,473 core genes confirmed the genetic coherenc of the U.S. *Csp* population (**Figure 6B**). Bootstrap support below 50 at the base of the U.S. *Csp* clade still leads to multifurcation confirming a recent radiation event from a common ancestor. However, compared to the 76-gene tree in **Figure 6A**, the U.S. *Csp* clade in the 1,473-gene tree has more resolution and a few subclades with high bootstrap support become evident. In most cases, these clades include isolates from different host species such as redbud, dogwood, and wax myrtle, indicating the absence of host adaptation. Since only one isolate from Indiana and one from Tennessee provided enough reads for high quality assemblies and all other isolates came from Virginia, nothing can be concluded about regional differentiation.

Since we only obtained high quality genome assemblies from 17 out of the 106 total samples, our ability to infer evolutionary relationships of U.S. *Csp* isolates using a core genome approach was limited. Thus, we also tried to use unassembled *Ceratobasidium* reads of a larger number of samples: the 17 samples that had provided high quality draft assemblies and 14 samples that had provided partial genome assemblies, which included two *Ct* samples from cacao from Indonesia. These reads were aligned against the *Ct* LOA1 genome, the genome of the closest available relative of the U.S. *Csp* lineage based on the above core genome analyses. 1,317,177 SNPs were identified and used for phylogenetic reconstruction (**Figure 7)**.

**Figure 7.**
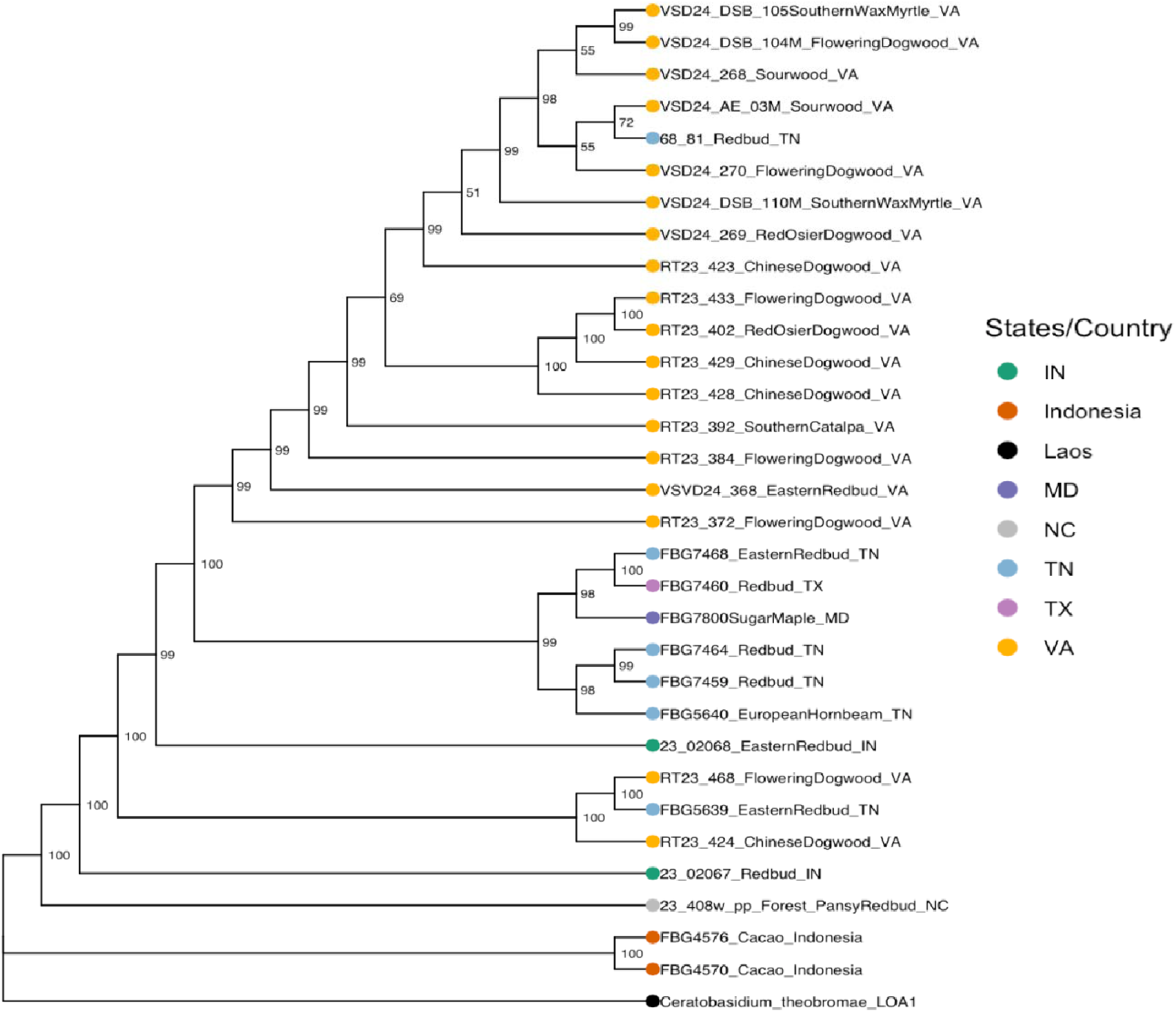
Maximum□likelihood phylogeny constructed under a `GTR+`Γ model from 32 *Ceratobasidium* isolates using 1.32 million SNPs identified by the NanoCaller tool by aligning reads against the *Ct* LOA1 genome, which was chosen as root. Ultrafast bootstrap values (≥ 50%) are shown at nodes.

The two Indonesian cacao isolates formed a separate clade with 100% bootstrap support, confirming the deep evolutionary split between U.S. *Csp* isolates and Southeast Asian *Ct* isolates seen in the two core genome trees (**Figure 6**). All U.S. *Csp* isolates form a single clade, also with 100% bootstrap support and thus also in agreement with the core genome trees. The only North Carolina isolate branched off from the most basal node of this clade, which is in lin with the epidemiological finding that trees with VSD symptoms were first observed in North Carolina in 2018, and *Csp* may have spread from there to the other U.S. states. Intriguingly, all but two Virginia isolates form a single well supported clade and the majority of Tennessee isolates form a second well supported clade suggesting that *Csp* populations in these two states may have started to genetically separate from each other. As already seen in the core genome tree, there are no host-specific lineages suggesting no adaptation to different host species.

### Exploring gene content differences between genomes of U.S. *Csp* isolates and other *Ceratobasidium* and *Rhizoctonia* isolates and among U.S. *Csp* isolates

All phylogenetic trees showed that the genomes of the U.S. *Csp* isolates form a single clade that is clearly distinct from the genomes of the Southeast Asian *Ct* isolates from cacao and cassava and all public genomes of other *Ceratobasidium* and *Rhizoctonia* isolates. However, just how different is this U.S. clade compared to all other isolates? To help answer this question, we performed a pangenome analysis. We found a total of 176,561 proteins in 12 *Ceratobasidium* and 3 *R. solani* genomes. This dataset incorporated the five highest quality individual genome assemblies of our U.S. samples, our comprehensive assembly, the genomes of the three *Ct* isolates from cacao and cassava, three public genomes of *Csp* isolates with the highest BUSCO scores, and the genomes of the three *R. solani* genomes used as outgroup in our core genome phylogeny (**Figure 6A**).

The pangenome analysis identified 10,913 non-redundant orthologous protein groups (orthogroups). Of these, 5,211 orthogroups (47.8%) were classified as core, being shared across every single analyzed genome. Additionally, 4,278 orthogroups (39.2%) were categorized as accessory, present in some but not all genomes, reflecting genomic variability within the species. Singletons, representing 13.1% of the orthogroups (1,424), consisted of genes unique to individual genomes, either due to differences in genome quality or representing true differences between individual isolates (**Figure 8A**). The gene-accumulation curve (**Figure 8B**) plots orthogroup counts against the number of genomes considered. Counts generally decline as *N* increases, with a pronounced spike in the rightmost bin (15 genomes) that reflects the core orthogroups shared by nearly all genomes, while the spread across intermediate bins indicates an open pangenome with substantial accessory variation among isolates.

**Figure 8.**
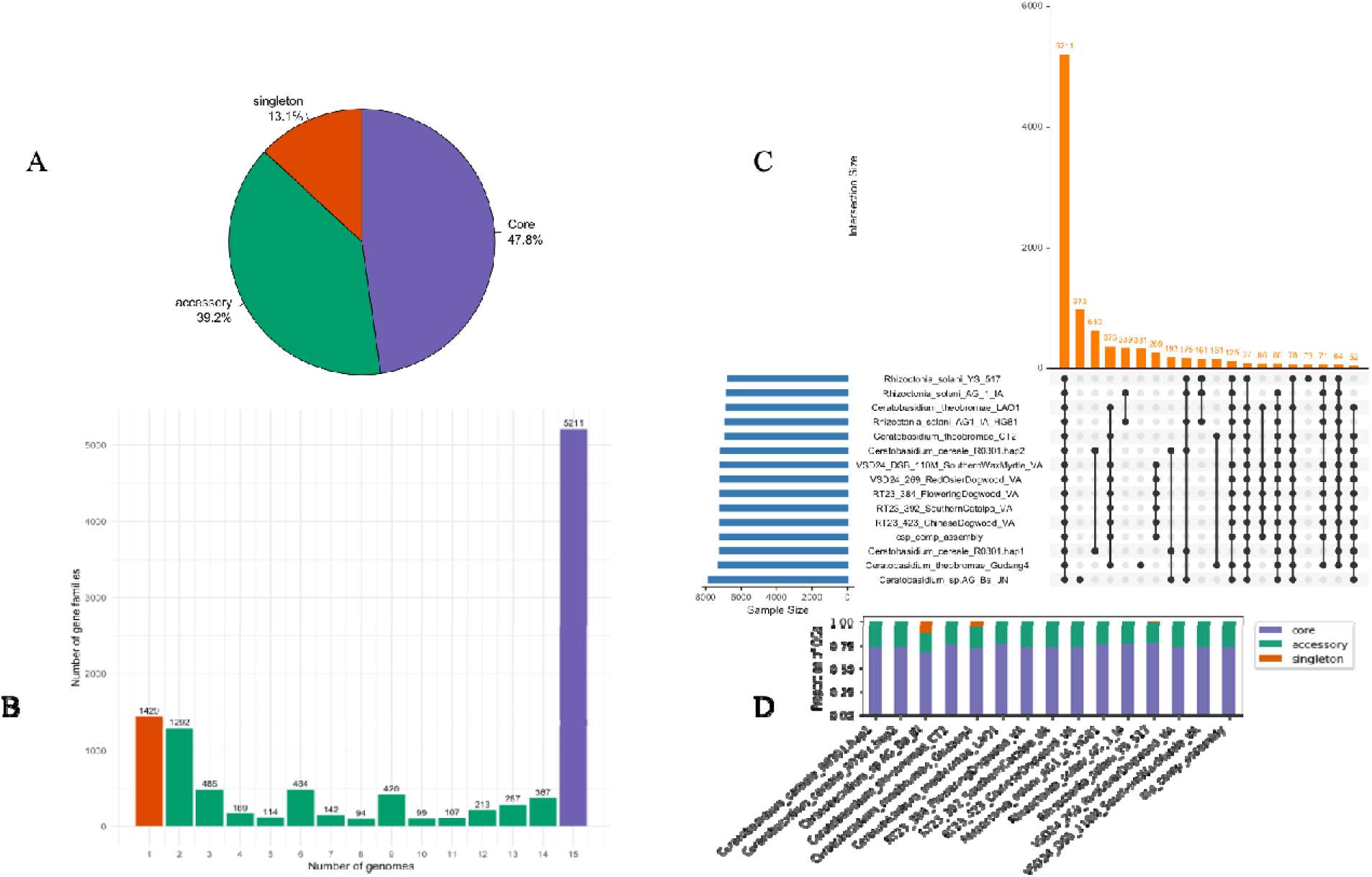
Pangenome analysis. **A.** Composition of core and accessory orthogroups and singleton genes in the *Ceratobasidium*/*Rhizoctonia* pan-genome. **B.** Gene accumulation curve (orthogroup count vs. number of genomes): each bar represents how many gene families are present when considering a specific number of genomes. The numbers decrease as more genomes are included, with a sharp increase near the rightmost bars (particularly at 15 genomes), reflecting the core gene families shared by nearly or all genomes. This curve illustrates how the pangenome grows and highlights both unique and shared gene content across genomes. **C**. UpSet plot showing intersections of orthogroups across 15 *Ceratobasidium* and *R. solani* genomes. Bar heights indicate the number of orthogroups for each presence/absence pattern (only intersections with ≥50 OGs are shown), with the connected dots below identifying the genomes included in each intersection. **D.** Composition of core, accessory and singleton gene per genome.

The UpSet plot (**Figure 8C**) shows how the accessory orthogroups are distributed. Importantly, 269 orthogroups were shared among the five genomes derived from US samples and the comprehensive assembly but absent from all other analyzed genomes, while 161 orthogroups were shared by the two *Ct* genomes from cacao but absent from all other analyzed genomes. This difference in gene content underscores the evolutionary separation between these two clades. An additional 86 genes were shared among the U.S. *Csp* genomes and the *Ct* LAO1 genome from cassava but absent from all other genomes while fewer than 50 genes were shared between the *Ct* LAO1 genome and the two *Ct* genomes from cacao or between the U.S. *Csp* genomes and the two *Ct* genomes from cacao. This further confirms the closer relationship between the U.S. *Csp* isolates with the *Ct* lineage from cassava than with the *Ct* genomes from cacao, which was also evident in the core genome tree in **Figure 6A**. Per-genome composition plots (**Figure 8D**) further indicate broadly similar proportions of core, accessory, and singleton content across isolates, with modest variation in singleton burden that likely reflects both assembly completeness and genuine lineage-specific gene gain/loss.

To hone in on the genomic diversity among the U.S. *Csp* genomes and their comparison with the *Ct* genomes, we conducted a second pangenome analysis of all 17 *Csp* genomes and 3 *Ct* genomes from the core-genome phylogeny (**Figure 6B**). We also included the comprehensive assembly because of the confidence we have in its completeness. Clustering 227,262 proteins from these 20 genomes resulted in 9,902 orthologous groups: 65.1% core, 31.7% accessory, and 3.2% singletons. The UpSet plot (**Figure S2**) reveals 6,446 core orthogroups found in all genomes. One can notice that each of the lowest quality genomes, such as *Csp* VSD24-268_Sourwood_VA or 23_02068_EasternRedbud_IN, are missing between 200 and 400 genes, probably because of their low genome completeness. Therefore, the true core genome of the analyzed genomes probably exceeds 7,000 orthogroups. Since we had observed two well-supported clades in **Figure 6B**, we were particularly curious to see if these two clades could be distinguished based on orthologue presence or absence, but no such trend is visible suggesting that the two clades have only diverged minimally so far.

As for the differences between the U.S. *Csp* genomes and the *Ct* genomes from Southeast Asia, this focused *Csp*/*Ct* pangenome analysis confirmed the substantial differences in orthogroup content found between the two groups in the wider pangenome analysis from **Figure 8**, only that the U.S. *Csp* genomes now shared slightly fewer orthogroups, 214 versus 269, probably because of the inclusion of the lower quality genomes with lower completeness.

### Prediction of effector genes in accessory genomes

For effector prediction, we targeted the accessory orthogroups identified in the 15 genomes used in the phylogenetic analysis (**Figure 6A**) and the pangenome analysis (**Figure 8)**. Accessory proteins per genome ranged from 2,184 in *Ct* LAO1 to 8,485 in *Csp* AG_Ba_Jn (**Table 2**). Out of the 16,294 accessory proteins, SignalP 6.0 predicted 1,431 to have N-terminal signal peptides. To exclude membrane-bound proteins from this group, we applied TMHMM v2.0c to remove sequences with at least one predicted transmembrane (TM) helix. After this filtering step, 1,231 proteins were retained. One of these proteins was removed because it contained a mitochondrial transit peptide identified by TargetP 2.0. None were found to have ER-retention motifs by ScanProsite (PS00014) or GPI-anchor signals by NetGPI 1.1. Therefore, 1230 secretome candidates were finally screened with EffectorP 3.0 in fungal mode, which partitioned them into effectors and non□effectors and further sub□classified effector candidates as apoplastic, cytoplasmic, or dual□localized (**Table 2**).

**Table 2.**
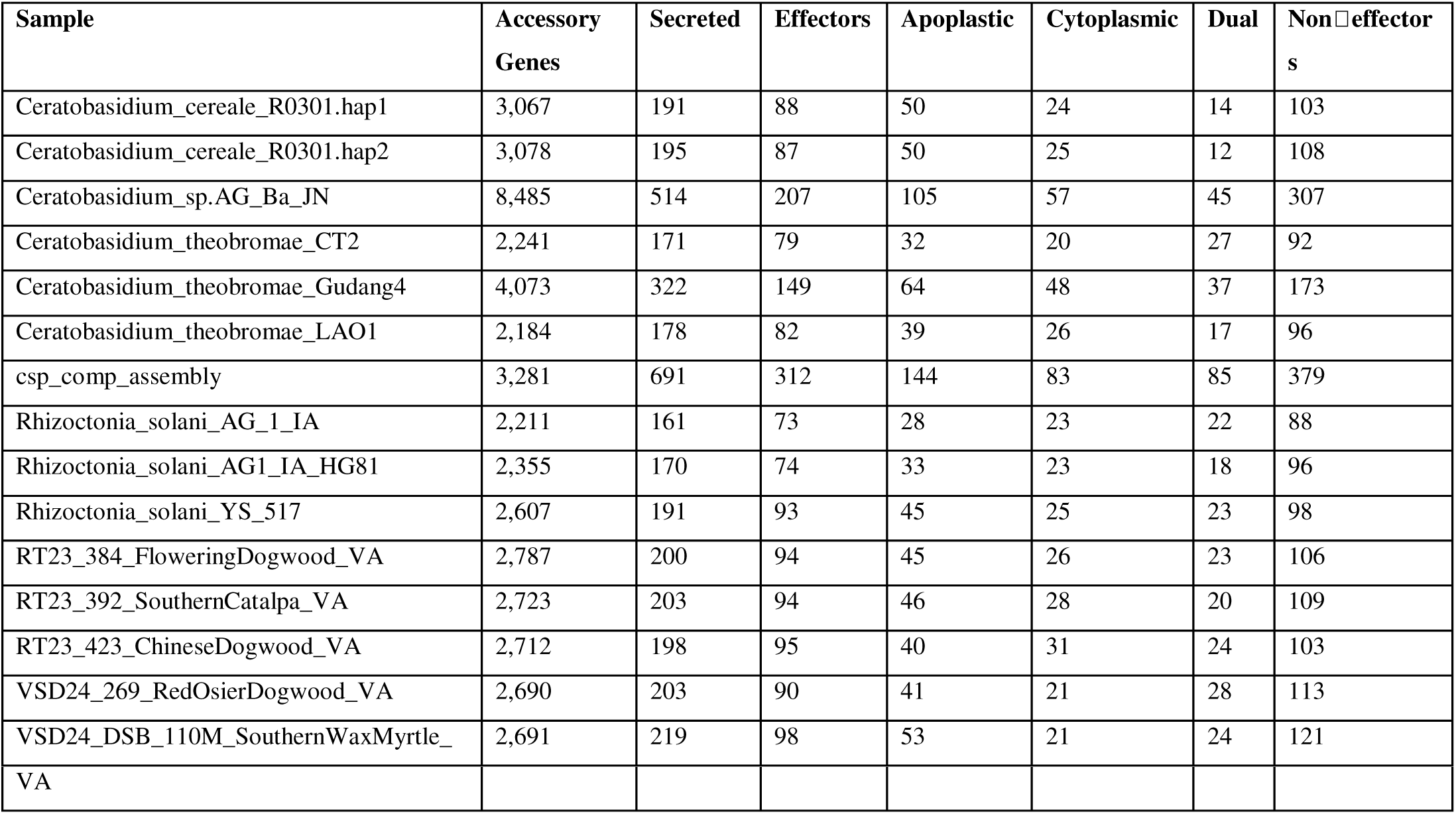
Summary of accessory gene content and effector predictions for *Ceratobasidium* and *R. solani* genomes.

To identify clade-specific effectors, we analyzed the predicted effector repertoires across the major clades identified in our core-genome phylogeny (**Figure 6A**). The non-U.S. *Csp* clade contained the highest number of unique predicted effectors, with 150 effectors specific to this group. The U.S. *Csp* clade harbored a distinct set of 34 unique effectors, while the *Ct* (cacao/cassava) clade had 52 unique effectors. The *R. solani* group displayed the lowest effector count, with only 10 identified effectors. These results reveal substantial differences in effector content between the US *Csp* clade and its sister clades, yielding a distinct set of effector candidates that can be further investigated for their potential roles in host specificity and virulence.

## Discussion

Since the first detection of *Csp* on an eastern redbud with VSD symptoms in Tennessee in 2019, VSD has quickly spread across 13 U.S. states and *Csp* has been detected molecularly on 50 genera of woody ornamentals (Liyanapathiranage et al., 2025; McClellan 2023). However, the fastidious nature of *Csp* has prevented Koch’s postulates to be completed and to establish *Csp* as the sole causative agent of VSD (Ali et al., 2019; Fredricks & Relman., 1996, Liyanapathiranage et al. 2024). Therefore, the first objective of this study was to determine if any other bacterial, fungal, or oomycete pathogen besides *Csp* is consistently associated with VSD and may contribute to the disease.

Metagenomic sequencing is a promising approach to answer this question since it does not require culturing and can potentially identify all microbes in a sample to the species rank and, if enough sequencing data are obtained, MAGs can be obtained and used in phylogenetic reconstruction (Asnicar et al., 2020; Czech et al., 2022). However, while metagenomics has become relatively established for the identification of bacterial pathogens, including plant pathogens (Duan et al., 2009; Piombo et al., 2021; Navgire et al., 2022), identification of fungi from metagenomic data is challenging and is still in its infancy (Donovan et al., 2018; Gökdemir et al., 2022; Avershina et al., 2025). One main issue is that taxonomic profiling of metagenomic sequences heavily relies on the availability of correctly classified reference genomes and only 659 fungal genomes are currently (July 2025) included in NCBI’s manually curated RefSeq database compared to 435,623 prokaryotic genomes (Goldfarb et al., 2025). Therefore, instead of using the standard RefSeq-based reference database provided by the developers of the taxonomic profiler Kraken2, we built a custom fungal and oomycete Kraken2 database using all genomes from NCBI’s Assembly database. However, since curation by NCBI of the Assembly database is limited, genomes may be misclassified, leading to errors in the Kraken2 database constructed from these genomes, and, ultimately, giving false negative and positive results. This is probably the reason that several fungal species identified in our metagenomes by Kraken2 were not confirmed when using other tools (sourmash and minimap2). The same problem of false positives even happened when using Kraken2 with a standard RefSeq-based database for identification of bacterial species, possibly a result of misclassified genomes even presents in NCBI’s RefSeq database (Bagheri et al., 2020; Chorlton SD., 2024; et al., 2024) and/or limitations inherent to the Kraken2 algorithm (Breitwieser et al., 2018; Wood et al., 2019; Lu et al., 2023). Therefore, improved Kraken2 reference databases are urgently needed, and it is key not to rely on a single taxonomic profiler when analyzing metagenomic data.

Importantly though, we were able to identify sequencing reads as *Csp* in all samples using both Kraken2 in combination with our custom fungal database (which included *Ct* genomes while the standard RefSeq-based Kraken2 database did not) and mapping to two *Ct* reference genomes using minimap2. We obtained a similar number of reads with both tools giving us confidence that the number of false positive and negative reads assigned to *Csp* by these tools were limited. However, independently of the bioinformatics tools, we cannot exclude that a small number of *Csp* sequences were the results of barcode hopping, whereby some DNA molecules from one sample get erroneously labeled with the barcode from another sample, as has been reported to occur at low frequency with nanopore sequencing (Ezpeleta et al., 2022).

Since no pathogen besides *Csp* was consistently detected in the plant samples, our results confirmed that no other pathogen beyond *Csp* plays a significant role in VSD. The presence of the canker and dieback pathogen *B. dothidea* in some samples suggests that this pathogen may occasionally contribute to the severity of VSD, but symptoms caused by *B. dothidea* were previously found to be distinct from VSD and the pathogen was excluded from being the VSD-causing agent (Liyanapathiranage et al., 2025). The bacterial species *Stenotrophomonas maltophilia*, sometimes reported as plant pathogen (Hu et al., 2021), was mainly identified in mycelium samples but not in plant samples and can thus also be excluded as causing VSD.

An unexpected result was the low number of *Csp* reads in many of the plant samples. We speculate that in some cases samples may have been taken from tissue with low pathogen abundance or late during infection (with *Csp* DNA already degraded by the time of sampling) or that the employed DNA extraction protocols were relatively inefficient in releasing DNA from *Csp* cells compared to releasing DNA from other fungal or bacterial cells. Moreover, metabolites present in woody tissue are known to interfere with DNA extraction (Rizzo et al., 2022). Thus, for sensitive detection of *Csp*, it is important to still improve DNA extraction protocols. Finally, it is important to point out that taxa for which closely related genomes have not been published (and are thus missing from our databases) may have been present but remained undetectable.

After we had confirmed *Csp* as the VSD pathogen with the metagenomic results, we attempted assembly of *Csp* genomes from as many samples as possible for downstream core genome phylogenetic and pangenome analyses. For including genomes in downstream analysis, we used a minimum completeness score threshold of 70%, based on the Agaricomycetes BUSCO dataset (Manni et al., 2021). This may seem a relatively low completeness score but even our comprehensive assembly with a genome coverage of 250X (*i.e.*, every region of the assembly was covered on average by 250 reads making it very unlikely that some genomic regions were not covered at all) only reached a completeness score of 83.7. This suggests that the *Ceratobasidium* genus may be missing some of the genes present in most of the other sequenced Agaricomycetes genomes or that some genomic regions either escaped nanopore sequencing or could not be assembled. However, since our average read length was just shy of 3,000bp and nanopore sequencing is particularly well suited for assembling structurally complex regions and repeat regions (Zheng et al., 2020; Wang et al., 2021; Kinkar et al., 2021), this seems unlikely. Also, since the pangenome analysis only identified 1 singleton gene in the comprehensive assembly and only between 0 and 38 singletons in the 12 best individual assemblies (which had completeness scores of 83-84% based on BUSCO agaricomycetes_odb10), we conclude that our best individual assemblies have higher completeness than what the BUSCO scores suggest. Moreover, the pangenome results suggest that the genome assemblies present very low levels of contamination with other microbial sequences or plant sequences, and that sequencing errors did not significantly affect gene calling. Finally, the highly similar gene content between the best individual *Csp* assemblies (assembled from reads that mapped to genomes of the related, yet distinct, Southeast Asian *Ct* lineage) and the comprehensive assembly (which was based on all non-plant reads from five samples) shows that when using long reads, assembling genomes from reads mapped to a related genome has a low risk of accidentally excluding genomic regions that are possibly absent from the related genome used for mapping.

Once the genome assemblies were obtained, we built one 76-gene core genome tree to investigate the evolutionary relationships between the *Csp* genomes from the U.S. and related *Ceratobasidium* and *Rhizoctonia* species from other world regions, and one 1,473-gene core genome tree to unravel the evolutionary relationships among the *Csp* genomes from the U.S. The results from the first tree mainly confirmed a previously published tree based on the ITS, LSU, ATP6, TEF1, and RPB2 genes (Liyanapathiranage et al., 2024). However, probably because of the higher number of loci used (76 compared to 5), a single node at the base of all *Ceratobasidium* genomes had high bootstrap support confirming that the *Ceratobasidium* genus is monophyletic while the 5-gene tree could not separate the *Ceratobasidium* genomes from the *Rhizoctonia* genomes. Moreover, the 76-gene tree subdivided the *Ceratobasidium* clade into two strongly supported subclades: one consisting of the Southeast Asian *Ct* genomes and the U.S. *Csp* genomes and the other consisting of all other *Ceratobasidium* species. Interestingly, both the 76-gene tree and the 1,473-gene tree provide strong support for the *Csp* genomes from the U.S. sharing a more recent common ancestor with the *Ct* LAO1 genome from cassava in Laos than with the *Ct* genomes from cacao in Indonesia. Because of the small number of genomes that were available, this may be a coincidence. However, the result aligns with the earlier finding that the ITS regions of *Csp* isolates in the U.S. are identical to ITS sequences of *Ceratobasidium* isolates from Japanese honeysuckle in China (Liyanapathiranage et al., 2024), of which Laos is a direct neighbor, further suggesting that the U.S. *Csp* lineage may have its origin in that geographic region.

Regarding the evolutionary relationships among U.S. isolates, the 76-gene tree did not provide support for any subclades. This is not surprising since their most recent common ancestor may have existed not much longer than 7 years ago when the first trees with VSD symptoms were observed. Therefore, few mutations may have accumulated in the 76 core genes since then. The 1,473-gene tree did reveal two well-supported individual clades, both including samples from several different plant genera suggesting no adaptation to different hosts. Because only two clades had strong bootstrap support and since most of our best assemblies were of samples collected in one geographic region of Virginia, we could not make any conclusions about possible dissemination pathways. Therefore, we attempted construction of a SNP-based phylogeny based on alignment of unassembled individual reads against the *Ct* LAO1 genome. Since assembly was not required, this allowed us to include many more samples. While ONT significantly reduced their sequencing error rate over the years, it is still around 1% (Delahaye et al., 2020; Sahlin et al., 2021). Probably because of this relatively high error rate, even when using the most stringent SNP-calling parameters, an excessively high number of 1.32 million SNPs were called. Random sequencing errors are expected to occur independently in each genome and are not expected to falsely group genomes together into clades. However, this high number of SNPs still reduced our confidence in the obtained SNP tree (even though bootstrap support for most clades was very high). Therefore, although the location of the only North Carolina sample at the very base of the U.S. clade (suggesting that it is most similar to the most recent common ancestor of all U.S. isolates) agrees with the finding that VSD was first observed in North Carolina and *Csp* may have spread to the rest of the U.S. from there, this result will need to be followed up on. Additional sampling and sequencing, ideally using technologies with lower error rates, such as Illumina or PacBio HiFi, should be performed. Similarly, the possible separation of *Csp* into a Virginia and a Tennessee sub-population also still need to be confirmed.

The final question we wanted to answer was: are there any signs in the genomes of the U.S. *Csp* isolates that suggest that the *Csp* lineage adapted to woody ornamentals and North American climate and, possibly, represents a new *Ceratobasidium* species distinct from *Ct*? Our pangenome and effector gene prediction analyses found substantial differences in gene content between the genomes in the U.S. *Csp* clade and genomes in its sister clades. The U.S. *Csp* clade contained 214 to 269 unique genes, depending on which genomes were compared. Of these, up to 34 genes were predicted to encode secreted *Csp*-specific effectors, which possibly contribute to virulence by suppressing plant immunity or degrading plant cell walls, typical mechanisms used by fungal plant pathogens (de Sain and Rep, 2015; Jashni et al., 2015). Unfortunately, following up on these findings with functional studies of pathogenicity and host specificity (Gil-Ordóñez et al., 2024; De La Fuente et al., 2022) will be complicated because of the fastidious nature of *Csp*, making in unlikely to unravel the basis of the exceptional wide host range of the U.S. *Csp* lineage any time soon.

Interestingly, the *Ct* lineage presented by the only available LAO1 genome from cassava in Laos is not only more closely related to the U.S. *Csp* lineage based on phylogeny but also shares more genes with the U.S. *Csp* lineage than the cacao lineage, further suggesting that the U.S. *Csp* lineage may have originated from a pathogen population located in geographic proximity to Laos. Therefore, the question about speciation is more complex than we anticipated. We may either have just one *Ct* species including *Ct* from cacao and from cassava and *Csp* from ornamentals in the U.S., two species (either grouping *Ct* from cassava in Laos with the U.S. *Csp* lineage or with the *Ct* lineage from cacao), or three separate species. Only a more in-depth taxonomic analysis including additional genome-sequenced isolates, and possibly phenotypic analyses, will be able to answer this question.

In conclusion, this study revealed how metagenomics using long read sequencing can overcome the challenges involved with the absence of pure cultures when dealing with fastidious fungal pathogens. While we encountered many issues with false positives during taxonomic profiling, the use of multiple, parallel approaches and a custom database still allowed us to confidently conclude that *Csp* is the only causative agent of VSD in U.S. ornamentals. Also, the long reads allowed for assembly of several high-quality draft genomes giving insight into the relationships between the U.S. *Csp* lineage and related pathogens. On the other hand, the still relatively high error rate of ONT reduced our confidence in the obtained SNP-based tree, which has the potential to give additional insights into pathogen dissemination patterns. Also, our high-quality genomes allowed us to perform a solid pangenome analysis providing the basis for a future taxonomic investigation, functional studies of pathogenicity and host range, and development of new molecular markers for highly specific, high throughput detection assays.

## Data Availability

The metagenomic sequencing data generated in this study have been deposited in the NCBI Sequence Read Archive (SRA) under BioProject accession number PRJNA1290818. A total of 106 sequence datasets is available, with associated BioSample metadata provided in the SRA submission. Genome assemblies for 17 isolates have been submitted to GenBank under BioProject PRJNA1338224. Custom scripts, pipelines, intermediate files, and databases used for analysis are publicly available in permanent Zenodo repositories: 1. core scripts, intermediate tables, and figure notebooks: https://doi.org/10.5281/zenodo.17344926, 2. single-nucleotide polymorphism (SNP) call files: https://doi.org/10.5281/zenodo.17315410, 3. Kraken2 classification outputs: https://doi.org/10.5281/zenodo.17220370 and 4. Hash (sourmash) database for fungi and oomycetes database: https://zenodo.org/records/17527097. Note that for the Kraken2 database, we only provided a combined_mapping.txt file in the scripts directory with list of genomes instead of the formatted database since the file size exceeded Zenodo’s file size limit.

## References

Abdelrazek S, Salamanca LR, Vinatzer BA. Metagenomic Sequencing of Tomato Plants with Wilt Symptoms Allows for Strain-Level Pathogen Identification and Genome-Based Characterization. Phytopathology. 2025 Apr;115(4):354–366. doi: 10.1094/PHYTO-09-24-0279-R. Epub 2025 Apr 25. PMID: 39752554.

Ali SS, Asman A, Shao J, Firmansyah AP, Susilo AW, Rosmana A, McMahon P, Junaid M, Guest D, Kheng TY, Meinhardt LW, Bailey BA. Draft genome sequence of fastidious pathogen *Ceratobasidium theobromae*, which causes vascular-streak dieback in *Theobroma cacao*. Fungal Biol Biotechnol. 2019 Sep 30;6:14. doi: 10.1186/s40694-019-0077-6. PMID: 31583107; PMCID: PMC6767637.

Almagro Armenteros JJ, Salvatore M, Emanuelsson O, Winther O, von Heijne G, Elofsson A, Nielsen H. Detecting sequence signals in targeting peptides using deep learning. Life Sci Alliance. 2019 Sep 30;2(5):e201900429. doi: 10.26508/lsa.201900429. PMID: 31570514; PMCID: PMC6769257.

Altschul SF, Gish W, Miller W, Myers EW, Lipman DJ. Basic local alignment search tool. J Mol Biol. 1990 Oct 5;215(3):403–10. doi: 10.1016/S0022-2836(05)80360-2. PMID: 2231712.

Asnicar, F., Thomas, A.M., Beghini, F. et al. Precise phylogenetic analysis of microbial isolates and genomes from metagenomes using PhyloPhlAn 3.0. Nat Commun 11, 2500 (2020). 10.1038/s41467-020-16366-7

Avershina, E., Qureshi, A.I., Winther-Larsen, H.C. et al. Challenges in capturing the mycobiome from shotgun metagenome data: lack of software and databases. Microbiome 13, 66 (2025). 10.1186/s40168-025-02048-3

Avin FA, Liyanapathiranage P, Bonkowski J, Bily D, Rodriguez Salamanca L, Olson J, Wallace S, McConnell M, Bec S, Baysal-Gurel F. Development of Molecular Markers for the Detection of *Ceratobasidium* sp. D.P. Rogers Associated with Vascular Streak Dieback of Woody Ornamentals in the United States. Plant Dis. 2025 May 30. doi: 10.1094/PDIS-01-25-0005-RE. Epub ahead of print. PMID: 40445165.

Bagheri H, Severin AJ, Rajan H. Detecting and correcting misclassified sequences in the large-scale public databases. Bioinformatics. 2020 Sep 15;36(18):4699–4705. doi: 10.1093/bioinformatics/btaa586. PMID: 32579213; PMCID: PMC7821992.

Banos, S., Lentendu, G., Kopf, A. et al. A comprehensive fungi-specific 18S rRNA gene sequence primer toolkit suited for diverse research issues and sequencing platforms. BMC Microbiol 18, 190 (2018). 10.1186/s12866-018-1331-4

Baysal-Gurel, F., and Liyanapathiranage, P. 2024. Vascular streak dieback is an emerging threat to the Redbud nursery production in the southeastern United States (ANR-PATH-1-2024). Tennessee State University Extension Publications. Retrieved March 8, 2024, from https://digitalscholarship.tnstate.edu/cgi/viewcontent.cgi?article=1180&context=extension

Bhunjun CS, Phillips AJL, Jayawardena RS, Promputtha I, Hyde KD. Importance of Molecular Data to Identify Fungal Plant Pathogens and Guidelines for Pathogenicity Testing Based on Koch’s Postulates. Pathogens. 2021 Aug 28;10(9):1096. doi: 10.3390/pathogens10091096. PMID: 34578129; PMCID: PMC8465164.

Bily D, Gyatso T, Avin FA, Bonkowski J, Liyanapathiranage P, Rodriguez Salamanca L, Vinatzer B, Baysal-Gurel F. A *Ceratobasidium* sp. D.P. Rogers associated with vascular streak dieback of woody ornamental plants in Virginia, U.S.A. Plant Dis. 2025 Jun 5. doi: 10.1094/PDIS-02-25-0375-RE. Epub ahead of print. PMID: 40471229.

Breitwieser, F.P., Baker, D.N. & Salzberg, S.L. KrakenUniq: confident and fast metagenomics classification using unique k-mer counts. Genome Biol 19, 198 (2018). 10.1186/s13059-018-1568-0

Brown, et al, (2016), sourmash: a library for MinHash sketching of DNA, Journal of Open-Source Software, 1(5), 27, doi:10.21105/joss.00027

Buchfink, B., Reuter, K. & Drost, HG. Sensitive protein alignments at tree-of-life scale using DIAMOND. Nat Methods 18, 366–368 (2021). 10.1038/s41592-021-01101-x

Capella-Gutiérrez S, Silla-Martínez JM, Gabaldón T. trimAl: a tool for automated alignment trimming in large-scale phylogenetic analyses. Bioinformatics. 2009 Aug 1;25(15):1972–3. doi: 10.1093/bioinformatics/btp348. Epub 2009 Jun 8. PMID: 19505945; PMCID: PMC2712344.

Català S, Pérez-Sierra A, Abad-Campos P. The use of genus-specific amplicon pyrosequencing to assess phytophthora species diversity using eDNA from soil and water in Northern Spain. PLoS One. 2015 Mar 16;10(3):e0119311. doi: 10.1371/journal.pone.0119311. PMID: 25775250; PMCID: PMC4361056.

Chklovski, A., Parks, D.H., Woodcroft, B.J. et al. CheckM2: a rapid, scalable and accurate tool for assessing microbial genome quality using machine learning. Nat Methods 20, 1203–1212 (2023). 10.1038/s41592-023-01940-w

Chorlton SD. Ten common issues with reference sequence databases and how to mitigate them. Front Bioinform. 2024 Mar 15;4:1278228. doi: 10.3389/fbinf.2024.1278228. PMID: 38560517; PMCID: PMC10978663.

Clarridge JE 3rd. Impact of 16S rRNA gene sequence analysis for identification of bacteria on clinical microbiology and infectious diseases. Clin Microbiol Rev. 2004 Oct;17(4):840–62, table of contents. doi: 10.1128/CMR.17.4.840-862.2004. PMID: 15489351; PMCID: PMC523561.

Csardi G, Nepusz T (2006). “The igraph software package for complex network research.” InterJournal, Complex Systems, 1695. https://igraph.org.

Czech L, Stamatakis A, Dunthorn M, Barbera P. Metagenomic Analysis Using Phylogenetic Placement-A Review of the First Decade. Front Bioinform. 2022 May 26;2:871393. doi: 10.3389/fbinf.2022.871393. PMID: 36304302; PMCID: PMC9580882.

De La Fuente, L., Merfa, M. V., Cobine, P. A. & Coleman, J. J. Pathogen adaptation to the xylem environment. Annu. Rev. Phytopathol. 60, 161–186 (2022).

Delahaye C, Nicolas J. Sequencing DNA with nanopores: Troubles and biases. PLoS One. 2021 Oct 1;16(10):e0257521. doi: 10.1371/journal.pone.0257521. PMID: 34597327; PMCID: PMC8486125.

Dinghua Li, Chi-Man Liu, Ruibang Luo, Kunihiko Sadakane, Tak-Wah Lam, MEGAHIT: an ultra-fast single-node solution for large and complex metagenomics assembly via succinct de Bruijn graph, Bioinformatics, Volume 31, Issue 10, May 2015, Pages 1674–1676, 10.1093/bioinformatics/btv033

Donovan PD, Gonzalez G, Higgins DG, Butler G, Ito K. Identification of fungi in shotgun metagenomics datasets. PLoS One. 2018 Feb 14;13(2):e0192898. doi: 10.1371/journal.pone.0192898. PMID: 29444186; PMCID: PMC5812651.

Duan Y, Zhou L, Hall DG, Li W, Doddapaneni H, Lin H, Liu L, Vahling CM, Gabriel DW, Williams KP, Dickerman A, Sun Y, Gottwald T. Complete genome sequence of citrus huanglongbing bacterium, *’Candidatus Liberibacter asiaticus*’ obtained through metagenomics. Mol Plant Microbe Interact. 2009 Aug;22(8):1011–20. doi: 10.1094/MPMI-22-8-1011. PMID: 19589076.

Edgar, R.C. MUSCLE: a multiple sequence alignment method with reduced time and space complexity. BMC Bioinformatics 5, 113 (2004). 10.1186/1471-2105-5-113

Edouard de Castro, Christian J. A. Sigrist, Alexandre Gattiker, Virginie Bulliard, Petra S. Langendijk-Genevaux, Elisabeth Gasteiger, Amos Bairoch, Nicolas Hulo, ScanProsite: detection of PROSITE signature matches and ProRule-associated functional and structural residues in proteins, Nucleic Acids Research, Volume 34, Issue suppl_2, 1 July 2006, Pages W362–W365, 10.1093/nar/gkl124

Ezpeleta J, Garcia Labari I, Villanova GV, Bulacio P, Lavista-Llanos S, Posner V, Krsticevic F, Arranz S, Tapia E. Robust and scalable barcoding for massively parallel long-read sequencing. Sci Rep. 2022 May 10;12(1):7619. doi: 10.1038/s41598-022-11656-0. PMID: 35538127; PMCID: PMC9090787.

Emms, D.M., Kelly, S. OrthoFinder: phylogenetic orthology inference for comparative genomics. Genome Biol 20, 238 (2019). 10.1186/s13059-019-1832-y

Flynn JM, Hubley R, Goubert C, Rosen J, Clark AG, Feschotte C, Smit AF. RepeatModeler2 for automated genomic discovery of transposable element families. Proc Natl Acad Sci U S A. 2020 Apr 28;117(17):9451–9457. doi: 10.1073/pnas.1921046117. Epub 2020 Apr 16. PMID: 32300014; PMCID: PMC7196820.

Fredricks DN, Relman DA. Sequence-based identification of microbial pathogens: a reconsideration of Koch’s postulates. Clin Microbiol Rev. 1996 Jan;9(1):18–33. doi: 10.1128/CMR.9.1.18. PMID: 8665474; PMCID: PMC172879.

Geneve, R. (1991). Eastern Redbud (Cercis canadensis L.) and Judas Tree (Cercis siliquastrum L.)., 142-151. 10.1007/978-3-662-13231-9_8.

Gil-Ordóñez, A., Pardo, J.M., Sheat, S. et al. Isolation, genome analysis and tissue localization of *Ceratobasidium theobromae*, a new encounter pathogen of cassava in Southeast Asia. Sci Rep 14, 18139 (2024). 10.1038/s41598-024-69061-8

Gíslason, M. H., Nielsen, H., Almagro Armenteros, J. J., & Johansen, A. R. (2021). Prediction of GPI-anchored proteins with pointer neural networks. Current Research in Biotechnology, 3, 6–13. 10.1016/j.crbiot.2021.01.001

Gökdemir FŞ, İşeri ÖD, Sharma A, Achar PN, Eyidoğan F. Metagenomics Next Generation Sequencing (mNGS): An Exciting Tool for Early and Accurate Diagnostic of Fungal Pathogens in Plants. J Fungi (Basel). 2022 Nov 13;8(11):1195. doi: 10.3390/jof8111195. PMID: 36422016; PMCID: PMC9699264.

Goldfarb T, Kodali VK, Pujar S, Brover V, Robbertse B, Farrell CM, Oh DH, Astashyn A, Ermolaeva O, Haddad D, Hlavina W, Hoffman J, Jackson JD, Joardar VS, Kristensen D, Masterson P, McGarvey KM, McVeigh R, Mozes E, Murphy MR, Schafer SS, Souvorov A, Spurrier B, Strope PK, Sun H, Vatsan AR, Wallin C, Webb D, Brister JR, Hatcher E, Kimchi A, Klimke W, Marchler-Bauer A, Pruitt KD, Thibaud-Nissen F, Murphy TD. NCBI RefSeq: reference sequence standards through 25 years of curation and annotation. Nucleic Acids Res. 2025 Jan 6;53(D1):D243–D257. doi: 10.1093/nar/gkae1038. PMID: 39526381; PMCID: PMC11701664.

Gónzalez D, Rodriguez-Carres M, Boekhout T, Stalpers J, Kuramae EE, Nakatani AK, Vilgalys R, Cubeta MA. Phylogenetic relationships of *Rhizoctonia* fungi within the Cantharellales. Fungal Biol. 2016 Apr;120(4):603–619. doi: 10.1016/j.funbio.2016.01.012. Epub 2016 Jan 29. PMID: 27020160; PMCID: PMC5013834.

Gonzalez Garcia, V., Portal Onco, M. A. & Rubio Susan, V. Review. Biology and systematics of the genus *Rhizoctonia*. *Span*. J. Agric. Res. 4(1), 55–79. 10.5424/sjar/2006041-178 (2006).

Grünwald, N. (2012). Novel insights into the emergence of pathogens: the case of chestnut blight. Molecular Ecology, 21. 10.1111/j.1365-294X.2012.05597.x.

Guest, D., & Keane, P. (2007). Vascular-Streak Dieback: A New Encounter Disease of Cacao in Papua New Guinea and Southeast Asia Caused by the Obligate Basidiomycete *Oncobasidium theobromae*. Phytopathology, 97 12, 1654–7. 10.1094/PHYTO-97-12-1654

Handelsman J. Metagenomics: application of genomics to uncultured microorganisms. Microbiol Mol Biol Rev. 2004 Dec;68(4):669–85. doi: 10.1128/MMBR.68.4.669-685.2004. PMID: 15590779; PMCID: PMC539003.

Horta MAC, Steenwyk JL, Mead ME, Dos Santos LHB, Zhao S, Gibbons JG, Marcet-Houben M, Gabaldón T, Rokas A, Goldman GH. Examination of Genome-Wide Ortholog Variation in Clinical and Environmental Isolates of the Fungal Pathogen *Aspergillus fumigatus*. mBio. 2022 Aug 30;13(4):e0151922. doi: 10.1128/mbio.01519-22. Epub 2022 Jun 29. PMID: 35766381; PMCID: PMC9426589.

Hu M, Li C, Xue Y, Hu A, Chen S, Chen Y, Lu G, Zhou X, Zhou J. Isolation, Characterization, and Genomic Investigation of a Phytopathogenic Strain of *Stenotrophomonas maltophilia*. Phytopathology. 2021 Nov;111(11):2088–2099. doi: 10.1094/PHYTO-11-20-0501-R. Epub 2021 Nov 11. PMID: 33759550.

Irber, et al. (2024). sourmash v4: A multitool to quickly search, compare, and analyze genomic and metagenomic data sets. Journal of Open-Source Software, 9(98), 6830. 10.21105/joss.06830

Jacques F, Bolivar P, Pietras K, Hammarlund EU. Roadmap to the study of gene and protein phylogeny and evolution-A practical guide. PLoS One. 2023 Feb 24;18(2):e0279597. doi: 10.1371/journal.pone.0279597. PMID: 36827278; PMCID: PMC9955684.

Jesus Lozano-Fernandez, A Practical Guide to Design and Assess a Phylogenomic Study, Genome Biology and Evolution, Volume 14, Issue 9, September 2022, evac129, 10.1093/gbe/evac129

Johnson MA, Liu H, Bush E, Sharma P, Yang S, Mazloom R, Heath LS, Nita M, Li S, Vinatzer BA. Investigating plant disease outbreaks with long-read metagenomics: sensitive detection and highly resolved phylogenetic reconstruction applied to Xylella fastidiosa. Microb Genom. 2022 May;8(5):mgen000822. doi: 10.1099/mgen.0.000822. PMID: 35584001; PMCID: PMC9465077.

Kang DD, Li F, Kirton E, Thomas A, Egan R, An H, Wang Z. MetaBAT 2: an adaptive binning algorithm for robust and efficient genome reconstruction from metagenome assemblies. PeerJ. 2019 Jul 26;7:e7359. doi: 10.7717/peerj.7359. PMID: 31388474; PMCID: PMC6662567.

Kazutaka Katoh, John Rozewicki, Kazunori D Yamada, MAFFT online service: multiple sequence alignment, interactive sequence choice and visualization, Briefings in Bioinformatics, Volume 20, Issue 4, July 2019, Pages 1160–1166, 10.1093/bib/bbx108

Keane P, Flentje N, Lamb K. Investigation of vascular-streak dieback of cacao in Papua New Guinea. Aust J Biol Sci. 1972;25:553–64.

Kinkar L, Gasser RB, Webster BL, Rollinson D, Littlewood DTJ, Chang BCH, Stroehlein AJ, Korhonen PK, Young ND. Nanopore Sequencing Resolves Elusive Long Tandem-Repeat Regions in Mitochondrial Genomes. Int J Mol Sci. 2021 Feb 11;22(4):1811. doi: 10.3390/ijms22041811. PMID: 33670420; PMCID: PMC7918261.

Kolmogorov, M., Yuan, J., Lin, Y. et al. Assembly of long, error-prone reads using repeat graphs. Nat Biotechnol 37, 540–546 (2019). 10.1038/s41587-019-0072-8

Krogh A, Larsson B, von Heijne G, Sonnhammer EL. Predicting transmembrane protein topology with a hidden Markov model: application to complete genomes. J Mol Biol. 2001 Jan 19;305(3):567–80. doi: 10.1006/jmbi.2000.4315. PMID: 11152613.

Langmead B. 2024. Kraken 2/Bracken RefSeq ‘PlusPF’ database, build k2_pluspfp_20240605 [data set]. Center for Computational Biology, Johns Hopkins University. Available at: https://benlangmead.github.io/aws-indexes/k2. Accessed 20 June 2024.

Lauderdale, D. (2023). New vascular streak dieback fact sheet. North Carolina Cooperative Extension.

Leiva AM, Pardo JM, Arinaitwe W, Newby J, Vongphachanh P, Chittarath K, Oeurn S, Thi Hang L, Gil-Ordóñez A, Rodriguez R, Cuellar WJ. *Ceratobasidium* sp. is associated with cassava witches’ broom disease, a re-emerging threat to cassava cultivation in Southeast Asia. Sci Rep. 2023 Dec 15;13(1):22500. doi: 10.1038/s41598-023-49735-5. PMID: 38110543; PMCID: PMC10728180.

Li H, Handsaker B, Wysoker A, Fennell T, Ruan J, Homer N, Marth G, Abecasis G, Durbin R; 1000 Genome Project Data Processing Subgroup. The Sequence Alignment/Map format and SAMtools. Bioinformatics. 2009 Aug 15;25(16):2078–9. doi: 10.1093/bioinformatics/btp352. Epub 2009 Jun 8. PMID: 19505943; PMCID: PMC2723002.

Li H. Minimap2: pairwise alignment for nucleotide sequences. Bioinformatics. 2018 Sep 15;34(18):3094–3100. doi: 10.1093/bioinformatics/bty191. PMID: 29750242; PMCID: PMC6137996.

Liyanage, K. H. E., Liyanapathiranage, P., & Baysal-Gurel, F. (2025). Investigating the Economic Impact of Emerging Vascular Streak Dieback Threat to Redbuds in the US Nursery Industry. HortScience, 60(7), 1244–1250. https://www.doi.org/10.21273/HORTSCI18589-25

Liyanapathiranage P, Avin FA, Bonkowski J, Beckerman JL, Munster M, Hadziabdic D, Trigiano RN, Baysal-Gurel F. Vascular Streak Dieback: A Novel Threat to Redbud and Other Woody Ornamental Production in the United States. Plant Dis. 2025 Mar 3:PDIS04240905FE. doi: 10.1094/PDIS-04-24-0905-FE. Epub ahead of print. PMID: 39115954.

Lu J, Rincon N, Wood DE, Breitwieser FP, Pockrandt C, Langmead B, Salzberg SL, Steinegger M. Metagenome analysis using the Kraken software suite. Nat Protoc. 2022 Dec;17(12):2815–2839. doi: 10.1038/s41596-022-00738-y. Epub 2022 Sep 28. Erratum in: Nat Protoc. 2024 Aug 29. doi: 10.1038/s41596-024-01064-1. PMID: 36171387; PMCID: PMC9725748.

Lukas Käll, Anders Krogh, Erik L.L. Sonnhammer, Advantages of combined transmembrane topology and signal peptide prediction—the Phobius web server, Nucleic Acids Research, Volume 35, Issue suppl_2, 1 July 2007, Pages W429–W432, 10.1093/nar/gkm256

Mande SS, Mohammed MH, Ghosh TS. Classification of metagenomic sequences: methods and challenges. Brief Bioinform. 2012 Nov;13(6):669–81. doi: 10.1093/bib/bbs054. Epub 2012 Sep 8. PMID: 22962338.

Manni M, Berkeley MR, Seppey M, Simão FA, Zdobnov EM. BUSCO Update: Novel and Streamlined Workflows along with Broader and Deeper Phylogenetic Coverage for Scoring of Eukaryotic, Prokaryotic, and Viral Genomes. Mol Biol Evol. 2021 Sep 27;38(10):4647–4654. doi: 10.1093/molbev/msab199. PMID: 34320186; PMCID: PMC8476166.

Manni M, Berkeley MR, Seppey M, Zdobnov EM. BUSCO: Assessing Genomic Data Quality and Beyond. Curr Protoc. 2021 Dec;1(12):e323. doi: 10.1002/cpz1.323. PMID: 34936221.

Marsberg A, Kemler M, Jami F, Nagel JH, Postma-Smidt A, Naidoo S, Wingfield MJ, Crous PW, Spatafora JW, Hesse CN, Robbertse B, Slippers B. Botryosphaeria dothidea: a latent pathogen of global importance to woody plant health. Mol Plant Pathol. 2017 May;18(4):477–488. doi: 10.1111/mpp.12495. Epub 2016 Dec 13. PMID: 27682468; PMCID: PMC6638292.

Mcclellan, M. 2023. The redbud problem. https://www.nurserymag.com/article/the-redbudproblem/

McMahon P, Purwantara A. Vascular streak dieback (*Ceratobasidium theobromae*): history and biology. In: Bailey BA, Meinhardt LW, editors. Cacao diseases: a history of old enemies and new encounters. New York: Springer International Publishing; 2016. p. 307–35.

Meyer, F., Fritz, A., Deng, ZL. et al. Critical Assessment of Metagenome Interpretation: the second round of challenges. Nat Methods 19, 429–440 (2022). 10.1038/s41592-022-01431-4

Moliszewska E, Maculewicz D, Stępniewska H. Characterization of three-nucleate Rhizoctonia AG-E based on their morphology and phylogeny. Sci Rep. 2023 Oct 13;13(1):17328. doi: 10.1038/s41598-023-44448-1. PMID: 37833315; PMCID: PMC10575891.

Navgire, G.S., Goel, N., Sawhney, G. et al. Analysis and Interpretation of metagenomics data: an approach. Biol Proced Online 24, 18 (2022). 10.1186/s12575-022-00179-7

Nguyen LT, Schmidt HA, von Haeseler A, Minh BQ. IQ-TREE: a fast and effective stochastic algorithm for estimating maximum-likelihood phylogenies. Mol Biol Evol. 2015 Jan;32(1):268–74. doi: 10.1093/molbev/msu300. Epub 2014 Nov 3. PMID: 25371430; PMCID: PMC4271533.

Nurk S, Meleshko D, Korobeynikov A, Pevzner PA. metaSPAdes: a new versatile metagenomic assembler. Genome Res. 2017 May;27(5):824–834. doi: 10.1101/gr.213959.116. Epub 2017 Mar 15. PMID: 28298430; PMCID: PMC5411777.

Paradis E, Schliep K. ape 5.0: an environment for modern phylogenetics and evolutionary analyses in R. Bioinformatics. 2019 Feb 1;35(3):526–528. doi: 10.1093/bioinformatics/bty633. PMID: 30016406.

Pedersen BS, Quinlan AR. Mosdepth: quick coverage calculation for genomes and exomes. Bioinformatics. 2018 Mar 1;34(5):867–868. doi: 10.1093/bioinformatics/btx699. PMID: 29096012; PMCID: PMC6030888.

Pertea G, Pertea M. GFF Utilities: GffRead and GffCompare. F1000Res. 2020 Apr 28;9:ISCB Comm J-304. doi: 10.12688/f1000research.23297.2. PMID: 32489650; PMCID: PMC7222033.

Piombo E, Abdelfattah A, Droby S, Wisniewski M, Spadaro D, Schena L. Metagenomics Approaches for the Detection and Surveillance of Emerging and Recurrent Plant Pathogens. Microorganisms. 2021 Jan 16;9(1):188. doi: 10.3390/microorganisms9010188. PMID: 33467169; PMCID: PMC7830299.

Pritchard, L., Sharma, P., Mazloom, R., Pierce, T., Irber, L., Harrington, B., Heath, L., Brown, C.T., Vinatzer, B., 2022. genomeRxiv: a microbial whole-genome database and diagnostic marker design resource for classification, identification, and data sharing. Access Microbiology, 4(5).

Poudel RS, Belay K, Nelson B Jr, Brueggeman R, Underwood W. Population and genome-wide association studies of Sclerotinia sclerotiorum isolates collected from diverse host plants throughout the United States. Front Microbiol. 2023 Sep 27;14:1251003. doi: 10.3389/fmicb.2023.1251003. PMID: 37829452; PMCID: PMC10566370.

Quevillon E, Silventoinen V, Pillai S, Harte N, Mulder N, Apweiler R, Lopez R. InterProScan: protein domains identifier. Nucleic Acids Res. 2005 Jul 1;33(Web Server issue):W116–20. doi: 10.1093/nar/gki442. PMID: 15980438; PMCID: PMC1160203.

Rasmussen DA, Grünwald NJ. Phylogeographic Approaches to Characterize the Emergence of Plant Pathogens. Phytopathology. 2021 Jan;111(1):68–77. doi: 10.1094/PHYTO-07-20-0319-FI. Epub 2020 Nov 21. PMID: 33021879.

Rizzo D, Da Lio D, Bartolini L, Francia C, Aronadio A, Luchi N, Campigli S, Marchi G, Rossi E. DNA Extraction Methods to Obtain High DNA Quality from Different Plant Tissues. Methods Mol Biol. 2022;2536:91–101. doi: 10.1007/978-1-0716-2517-0_5. PMID: 35819599.

Roman-Reyna V and Crandall SG (2024) Seeing in the dark: a metagenomic approach can illuminate the drivers of plant disease. Front. Plant Sci. 15:1405042. doi: 10.3389/fpls.2024.1405042

Sahlin, K., Medvedev, P. Error correction enables use of Oxford Nanopore technology for reference-free transcriptome analysis. Nat Commun 12, 2 (2021). 10.1038/s41467-020-20340-8

Samuel T N Aroney, Rhys J P Newell, Jakob N Nissen, Antonio Pedro Camargo, Gene W Tyson, Ben J Woodcroft, CoverM: read alignment statistics for metagenomics, *Bioinformatics*, Volume 41, Issue 4, April 2025, btaf147, 10.1093/bioinformatics/btaf147

Samuels GJ, Ismaiel A, Rosmana A, Junaid M, Guest D, McMahon P, Keane P, Purwantara A, Lambert S, Rodriguez-Carres M, Cubeta MA. Vascular Streak Dieback of cacao in Southeast Asia and Melanesia: in planta detection of the pathogen and a new taxonomy. Fungal Biol. 2012 Jan;116(1):11–23. doi: 10.1016/j.funbio.2011.07.009. Epub 2011 Jul 23. PMID: 22208598.

Shen, W., Le, S., Li, Y., & Hu, F. (2016). SeqKit: A Cross-Platform and Ultrafast Toolkit for FASTA/Q File Manipulation. PLoS ONE, 11(10), e0163962. 10.1371/journal.pone.0163962

Skousen, J., Monteleone, A., Tyree, M., Swab, R., Groninger, J., Adams, M., Buckley, D., Wood, Smit AF. Repeat-Masker Open-3.0. 2004. [Google Scholar]

Sperschneider, J. & Dodds, P. N. EffectorP 3.0: Prediction of apoplastic and cytoplasmic effectors in fungi and oomycetes. Mol. Plant-Microb. Interact. 35(2), 146–156 (2022).

Stanke M, Keller O, Gunduz I, Hayes A, Waack S, Morgenstern B. AUGUSTUS: ab initio prediction of alternative transcripts. Nucleic Acids Res. 2006 Jul 1;34(Web Server issue):W435–9. doi: 10.1093/nar/gkl200. PMID: 16845043; PMCID: PMC1538822.

Sun K, Liu Y, Zhou X, Yin C, Zhang P, Yang Q, Mao L, Shentu X, Yu X. Nanopore sequencing technology and its application in plant virus diagnostics. Front Microbiol. 2022 Jul 25;13:939666. doi: 10.3389/fmicb.2022.939666. PMID: 35958160; PMCID: PMC9358452.

Tang B, Hu X, Yue M. Contaminated bacterial genome data in the public domains: Evidence and solution. J Infect. 2024 Dec;89(6):106369. doi: 10.1016/j.jinf.2024.106369. Epub 2024 Nov 26. PMID: 39608578.

Teufel F, Almagro Armenteros JJ, Johansen AR, Gíslason MH, Pihl SI, Tsirigos KD, Winther O, Brunak S, von Heijne G, Nielsen H. SignalP 6.0 predicts all five types of signal peptides using protein language models. Nat Biotechnol. 2022 Jul;40(7):1023–1025. doi: 10.1038/s41587-021-01156-3. Epub 2022 Jan 3. PMID: 34980915; PMCID: PMC9287161.

Tian L, Huang C, Mazloom R, Heath LS, Vinatzer BA. LINbase: a web server for genome-based identification of prokaryotes as members of crowdsourced taxa. Nucleic Acids Res. 2020 Jul 2;48(W1):W529–W537. doi: 10.1093/nar/gkaa190. PMID: 32232369; PMCID: PMC7319462.

Wang, Y., Zhao, Y., Bollas, A. et al. Nanopore sequencing technology, bioinformatics and applications. Nat Biotechnol 39, 1348–1365 (2021). 10.1038/s41587-021-01108-x

Wood, D.E., Lu, J. & Langmead, B. Improved metagenomic analysis with Kraken 2. Genome Biol 20, 257 (2019). 10.1186/s13059-019-1891-0

Woolhouse ME, Gowtage-Sequeria S. Host range and emerging and reemerging pathogens. Emerg Infect Dis. 2005 Dec;11(12):1842–7. doi: 10.3201/eid1112.050997. PMID: 16485468; PMCID: PMC3367654.):68-77. doi: 10.1094/PHYTO-07-20-0319-FI. Epub 2020 Nov 21. PMID: 33021879.

Wouter De Coster, Svenn D’Hert, Darrin T Schultz, Marc Cruts, Christine Van Broeckhoven, NanoPack: visualizing and processing long-read sequencing data, Bioinformatics, Volume 34, Issue 15, August 2018, Pages 2666–2669, 10.1093/bioinformatics/bty149

Yu G, Smith D, Zhu H, Guan Y, Lam TT (2017). “ggtree: an R package for visualization and annotation of phylogenetic trees with their covariates and other associated data.” Methods in Ecology and Evolution, 8, 28–36. doi:10.1111/2041-210X.12628

Zheng P, Zhou C, Ding Y, Liu B, Lu L, Zhu F, Duan S. Nanopore sequencing technology and its applications. MedComm (2020). 2023 Jul 10;4(4):e316. doi: 10.1002/mco2.316. PMID: 37441463; PMCID: PMC10333861.

