## Supplementary Tables and Figures for "Genomic insights into *Ceratobasidium* sp. associated with vascular streak dieback of woody ornamentals in the United States using a metagenomic sequencing approach": Supplementary Figures.docx (from another version of this preprint: version 2, 2025-09-09)

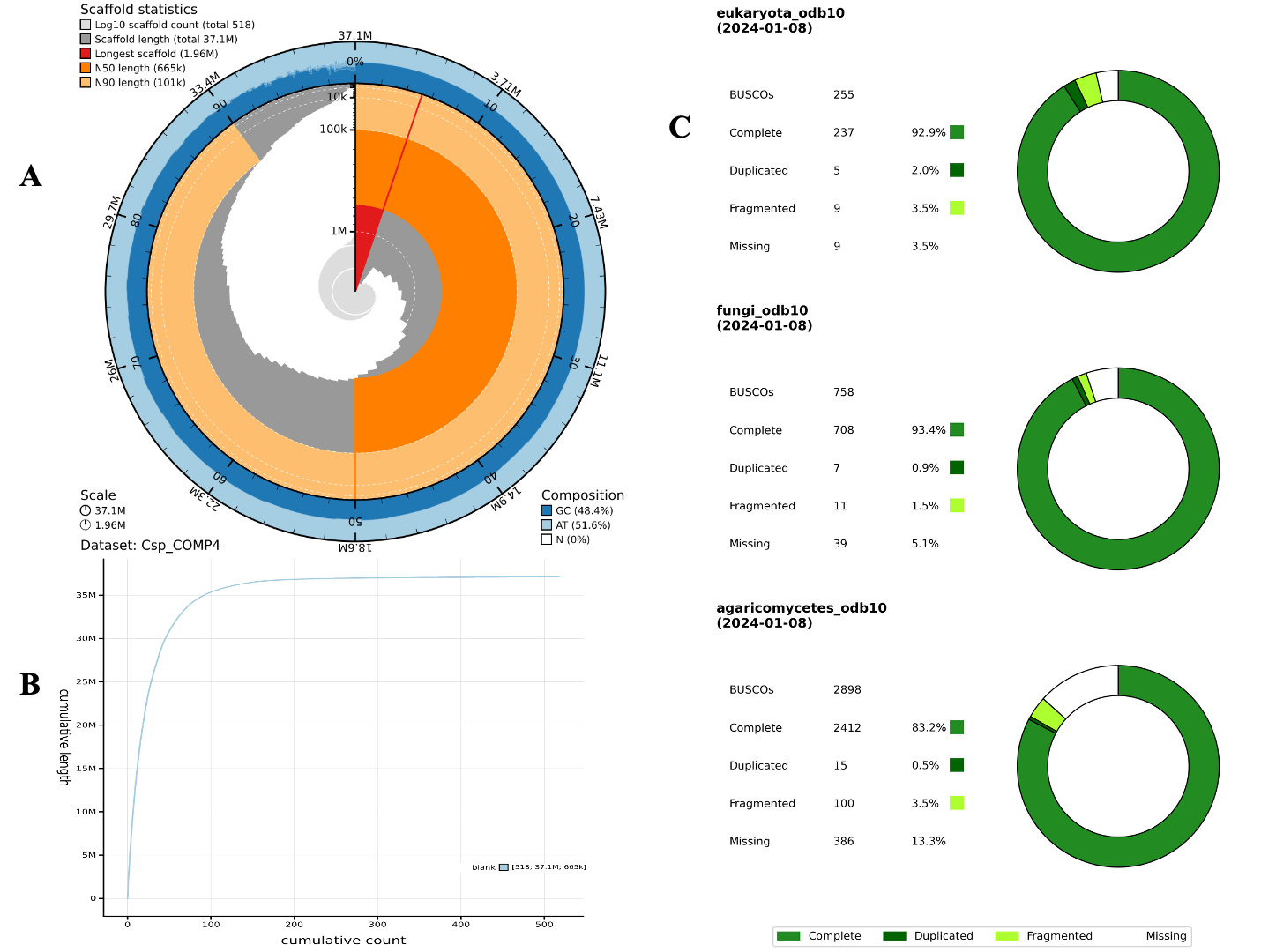

**Supplementary Figure 1. Genome assembly statistics and gene space completeness for the *Csp* comprehensive assembly.**

(**A**) Snail plot visualizing scaffold statistics for the comprehensive assembly. The assembly contains 518 scaffolds with a total length of 37.1 Mb. Scaffold lengths are arranged in decreasing order around the circumference, with the inner histogram (gray) plotted on a log₁₀ scale. The longest scaffold (1.96 Mb) is highlighted in red, the N50 scaffold length (665 kb) in orange, and the N90 (101 kb) in pale orange. Concentric rings indicate base composition, with GC content (48.4%) shown in blue and AT content (51.6%) in gray.

(**B**) Cumulative scaffold length plot, showing how cumulative assembly length (y-axis) increases with the number of scaffolds (x-axis), reaching 37.1 Mb at 518 scaffolds.

(**C**) BUSCO completeness assessment for the comprehensive assembly using three reference lineage datasets. For **eukaryota_odb10** (255 BUSCOs): 92.9% complete, 2.0% duplicated, 3.5% fragmented, 3.5% missing. For **fungi_odb10** (758 BUSCOs): 93.4% complete, 0.9% duplicated, 1.5% fragmented, 5.1% missing. For **agaricomycetes_odb10** (2,898 BUSCOs): 83.2% complete, 0.5% duplicated, 3.5% fragmented, 13.3% missing.

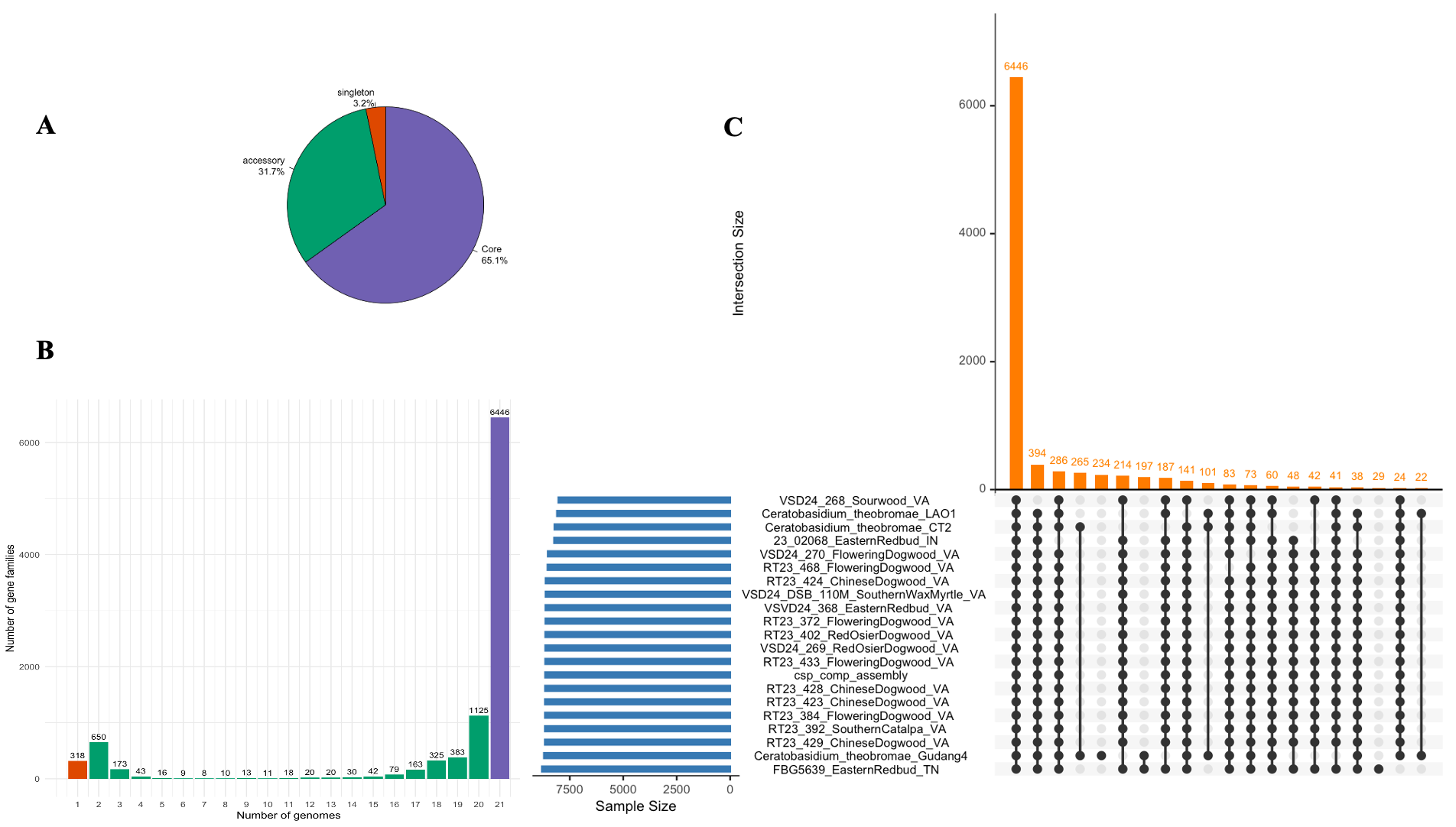

**Supplementary Figure 2.** Pangenome analysis of 17 US *Csp* genomes, the comprehensive *Csp* assembly, and three *Ct* reference genomes. **A.** Composition of core, accessory and singleton genes in the *Csp* and *Ct* pan-genome. **B.** Gene accumulation curve (orthogroup count vs. number of genomes): each bar represents how many gene families are present when considering a specific number of genomes. The numbers decrease as more genomes are included, with a sharp increase near the rightmost bars, reflecting the core gene families shared by nearly or all genomes. This curve illustrates how the pangenome grows and highlights both unique and shared gene content across genomes. **C**. UpSet plot showing intersections of orthogroups across all 21 genomes. Bar heights indicate the number of orthogroups for each presence/absence pattern, with the connected dots below identifying the genomes included in each intersection.
